# Modification of Non-photochemical Quenching Pathways in the C_4_ Model Plant *Setaria viridis* Revealed Shared and Unique Photoprotection Mechanisms as Compared to C_3_ Plants

**DOI:** 10.1101/2025.01.12.632622

**Authors:** Grace Milburn, Cheyenne M. Morris, Eileen Kosola, Dhruv Patel-Tupper, Jian Liu, Dominique H. Pham, Lucia Acosta-Gamboa, William D. Stone, Sarah Pardi, Kylee Hillman, William E. McHargue, Eric Becker, Xiaojun Kang, Josh Sumner, Catherine Bailey, Peter M. Thielen, Georg Jander, Cade N. Kane, Scott A. M. McAdam, Thomas J. Lawton, Dmitri A. Nusinow, Feng Zhang, Michael A. Gore, Jianlin Cheng, Krishna K. Niyogi, Ru Zhang

## Abstract

Light is essential for photosynthesis; however, excess light can increase the accumulation of photoinhibitory reactive oxygen species that reduce photosynthetic efficiency. Plants have evolved photoprotective non-photochemical quenching (NPQ) pathways to dissipate excess light energy. In tobacco and soybean (C_3_ plants), overexpression of three NPQ genes, *violaxanthin de-epoxidas*e (<u>V</u>DE), *Photosystem II Subunit S* (<u>P</u>sbS), and *zeaxanthin epoxidase* (<u>Z</u>EP), hereafter VPZ, resulted in faster NPQ induction and relaxation kinetics, and increased crop yields in field conditions. NPQ is well-studied in C_3_ plants; however, NPQ and the translatability of the VPZ approach in C_4_ plants is poorly understood. The green foxtail *Setaria viridis* is an excellent model to study photosynthesis and photoprotection in C_4_ plants. To understand the regulation of NPQ and photosynthesis in C_4_ plants, we performed transient overexpression in Setaria protoplasts and generated (and employed) stable transgenic Setaria plants overexpressing one of the three Arabidopsis NPQ genes or all three NPQ genes (AtVPZ lines). Overexpressing (OE) *AtVDE* and *AtZEP* in Setaria produced similar results as in C_3_ plants, with increased or reduced zeaxanthin (thus NPQ), respectively. However, overexpressing *AtPsbS* appeared to be challenging in Setaria, with largely reduced NPQ in protoplasts and under-represented homozygous AtPsbS-OE lines, potentially due to competitive and tight heterodimerization of AtPsbS and SvPsbS proteins. Furthermore, Setaria AtVPZ lines had increased zeaxanthin, faster NPQ induction, higher NPQ level, but slower NPQ relaxation. Despite this, AtVPZ lines had improved growth as compared to wildtype under several conditions, especially high temperatures, which is not related to the faster relaxation of NPQ but may be attributable to increased zeaxanthin and NPQ in C_4_ plants. Our results identified shared and unique characteristics of the NPQ pathway in C_4_ model Setaria as compared to C_3_ plants and provide insights to improve C_4_ crop yields under fluctuating environmental conditions.

## Introduction

Photosynthesis is the key driver for bioenergy and biomass production (Ort *et al*., 2015; Orr *et al*., 2017; Araus *et al*., 2021; Croce *et al*., 2024). It crucially depends on light; however, excess light damages photosynthesis, reducing plant growth and crop yield (Müller *et al*., 2001; Dietz, 2015; Pinnola & Bassi, 2018). Under optimized conditions, photosynthetic rates increase linearly at low light irradiances until saturation is reached at intensities often well below peak sunlight (Erickson *et al*., 2015; Pinnola & Bassi, 2018). The excess light plants absorb but cannot use for photosynthesis needs to be dissipated safely and efficiently via photoprotective pathways before damage occurs (Murchie & Ruban, 2020). Otherwise, excited chlorophyll that cannot transfer the energy for photosynthesis will interact with O_2_ to produce reactive oxygen species (ROS), which can damage chloroplastic lipids, membranes, and proteins (Mullineaux & Karpinski, 2002; Takahashi, 2011; Dietz, 2015). Under stressful conditions, e.g., high temperatures, drought, pathogen infection and others, photosynthesis saturates at a lower light intensity than under ideal conditions due to stress-induced inhibition of photosynthesis, making photoprotection even more important (Zhang & Sharkey, 2009; Sharkey & Zhang, 2010; Wang *et al*., 2018; Lu & Yao, 2018; Anderson *et al*., 2021).

Plants have several photoprotection pathways, one of the most important being a suite of non-photochemical quenching (NPQ) mechanisms (Müller *et al*., 2001; Rochaix, 2014; Pinnola & Bassi, 2018). NPQ has several components and its fastest and most dominant component is the energy-dependent quenching (qE), which is mainly modulated by three proteins: violaxanthin de-epoxidase (VDE), zeaxanthin epoxidase (ZEP), and the photosystem II (PSII) polypeptide Subunit S (PsbS) (Rochaix, 2014; Dietz, 2015; Ruban, 2016). Light triggers photosynthetic electron transport along the thylakoid membranes and proton translocation across the thylakoid membranes, acidifying the thylakoid lumen (Kramer *et al*., 2004; Baker *et al*., 2007). PsbS senses the change of lumen pH via two key protonatable glutamate residues, resulting in a conformational change which likely drives changes in thylakoid inter-protein interactions that transduce this signal to initiate qE via a yet unresolved mechanism (Li *et al*., 2004; Correa-Galvis *et al*., 2016; Krishnan-Schmieden *et al*., 2021; Chiariello *et al*., 2023; Marulanda Valencia & Pandit, 2024). VDE and ZEP are xanthophyll cycle enzymes: VDE is similarly activated by an acidic lumen pH, converting violaxanthin to intermediate antheraxanthin, and then to zeaxanthin; ZEP reverses the cycle, converting zeaxanthin back to violaxanthin (Jahns *et al*., 2009). Zeaxanthin has an important role in NPQ: increased zeaxanthin through VDE overexpression increases the induction rate of NPQ; whereas reduced zeaxanthin through ZEP overexpression decreases NPQ induction and amplitude (Hieber *et al*., 2001; Leonelli *et al*., 2016).

NPQ needs tight regulation to enable sufficient photoprotection without a cost to efficient photosynthesis. During the transition from high to low light, it is beneficial to relax NPQ quickly to maximize available light for photosynthesis (Zhu *et al*., 2010; Ghosh *et al*., 2023). In field conditions, sunlight often fluctuates between high to low light in several seconds, due to clouds passing by, and shading or movement of other leaves or plants. To improve photosynthesis in field conditions, a VPZ strategy was developed by modulating NPQ and overexpressing all three components of NPQ from the C_3_ model plant *Arabidopsis thaliana*: <u>V</u>DE, <u>P</u>sbS, and <u>Z</u>EP (Kromdijk *et al*., 2016; De Souza *et al*., 2022). These transgenic lines in tobacco and soybean plants had accelerated NPQ induction and also relaxation and improved yield in field conditions (Kromdijk *et al*., 2016; De Souza *et al*., 2022). However, the effects of the VPZ strategy have been difficult to reproduce in different plant species as it reduced plant growth and fitness in Arabidopsis and potato plants (Garcia-Molina & Leister, 2020; Lehretz *et al*., 2022), suggesting there are still many unknowns in translating optimized photoprotection in different plant species.

NPQ has been extensively studied in C_3_ plants, but much less so in C_4_ plants (Anderson *et al*., 2021). C_3_ plants use C_3_ photosynthesis, in which the first carbon compound produced contains three carbon atoms, e.g., rice and wheat (Yamori *et al*., 2014). Several important staple crops are C_4_ plants and utilize C_4_ photosynthesis, in which the first carbon compound produced contains four carbon atoms, e.g., maize, sorghum, and sugarcane. C_3_ photosynthesis uses one cell type for photosynthesis, mesophyll (M) cells, whereas C_4_ photosynthesis uses two cell types for photosynthesis, M and the adjacent bundle-sheath (BS) cells (Wang *et al*., 2011; von Caemmerer & Furbank, 2016). In C_4_ photosynthesis, CO_2_ is initially fixed in M cells to C_4_ acids, which are transported to BS cells to release CO_2_ around Ribulose-1,5-bisphosphate carboxylase/oxygenase (Rubisco) for carbon fixation, generating C_3_ acids (Sage, 2004; Wang *et al*., 2011; von Caemmerer & Furbank, 2016). This unique positioning of M and BS cells allows C_4_ plants to concentrate CO_2_ up to 10 X higher than ambient CO_2_ concentration around Rubisco, functioning as a carbon concentrating mechanism (CCM) (Danila *et al*., 2019). While this CCM comes at a bioenergetic cost to the plant, C_4_ photosynthesis is more efficient than C_3_ photosynthesis in hot and dry environments (Sage, 2004; Sage *et al*., 2012). While only 3% of the world’s terrestrial plant species use C_4_ photosynthesis, C_4_ plants are responsible for 20% of global gross primary productivity (Sage *et al*., 2012; Way *et al*., 2014). Despite the importance of C_4_ photosynthesis, its regulation, especially related to NPQ, is under-studied. C_4_ plants represent a unique platform to study the regulation of NPQ and learn how plants deal with the dilemma of photoprotection and light harvesting, particularly in stressful environments.

The C_4_ green foxtail grass, *Setaria viridis*, is an excellent model for studying the regulation of C_4_ photosynthesis and NPQ (Brutnell *et al*., 2010; Li & Brutnell, 2011). It has short stature and relatively short generation time (8∼10 weeks from seed to seed, 2 weeks from sowing to sufficient size for photosynthetic measurements). Setaria has a smaller genome size (400 Mb versus 2.3-2.7 Gb in maize) despite a similar number of genes as maize (38,000 in Setaria versus 33,000 in maize) (Mamidi *et al*., 2020; Thielen *et al*., 2020). It self-pollinates and produces hundreds of seeds (Brutnell *et al*., 2010; Li & Brutnell, 2011). Additionally, Setaria has highly efficient transformation protocols (from transformation to the T_0_ plantlets in about 10 weeks, with up to 25% transformation frequency) (Van Eck, 2018), advanced forward genetics protocols, and high-throughput phenotyping tools (Huang *et al*., 2016). More importantly, Setaria is an excellent model species for bioenergy crops, e.g., maize and sorghum, as all three belong to the same C_4_ photosynthesis subtype (NADP-ME type) (Wang *et al*., 2011; von Caemmerer & Furbank, 2016; Huang *et al*., 2016).

In this work, to investigate the regulation of NPQ in C_4_ plants, we first overexpressed Arabidopsis NPQ genes in Setaria protoplasts and revealed the surprising suppression effects on NPQ by overexpressing *AtPsbS* in Setaria protoplasts. We then generated stable transgenic Setaria plants by overexpressing one of the three Arabidopsis NPQ genes (AtNPQ lines, or AtVDE, AtZEP, AtPsbS, respectively) and employed Setaria transgenic lines overexpressing all three NPQ genes (AtVPZ lines) (Stone *et al*., 2024), performed thorough photosynthetic measurements in these plants, and phenotyped them under different environmental conditions. Our results show that AtVDE and AtZEP proteins worked similarly in C_4_ plants as in C_3_ plants; however, AtPsbS may be incompatible in Setaria, due to possible tighter binding to SvPsbS. Furthermore, the Setaria AtVPZ lines grew better than wildtype (WT) under several conditions, especially high temperature conditions, which is not related to the faster relaxation of NPQ but may be attributable to increased zeaxanthin and NPQ.

## Materials and Methods

### Plant growth conditions

*Setaria* WT and homozygous transgenic plants (T_3_-T_5_ lines) were grown under the control condition, 50% humidity, 31°C, plant-level light intensity of 250 μmol photons m^−2^ s^−1^, and 12/12-hour day/night cycle. Seeds were sown in Pro-Line C/V (Jolly Gardener, #18-10651). Plants were fertilized with 15-5-15 CA-MG LX (Jack’s Professional, #77940) and 15-16-17 Peat-Lite (Jack’s Professional, #77220). Seedlings in 5×5.5×6 cm pots were transplanted seven days after sowing (DAS) into 8×8×6 cm pots. Plants were watered morning and afternoon as needed. At 14 DAS, the fourth fully expanded leaf of a plant was used for photosynthetic measurements, tissue collection for RNA, protein, pigment, and abscisic acid (ABA) measurements.

### Protoplast isolation, transformation, and chlorophyll fluorescence measurements

Protoplasts were isolated using the third leaf of 12-day-old *Setaria viridis* WT plants grown in the control condition mentioned above; about ∼0.75 g with 15-20 leaves and the middle leaf segments (one centimeter from the base and tip of the leaf) were used for isolation. Leaves were cut vertically into small strips about one millimeter wide using a razor blade in a sterile petri dish in a drop of enzyme solution (0.3 g cellulase R10, 0.1 g macerozyme R10, 1 mL 0.2 M 4-morpholineethanesulfonic acid (MES), 10 mL 0.8 M mannitol, 20 μL 1 M MgCl_2,_ 20 μL 1 M CaCl_2,_ 7 μL 2-mercaptoethanol, 200 μL 10% bovine serum albumin, 100 μL 10 mg mL^-1^ carbenicillin, and 8.65 mL water). Then leaf pieces were transferred to a petri dish containing 20 mL of the enzyme solution mentioned above. The petri dish was loosely covered in foil and placed in a desiccator with a dark blanket and vacuum was applied for 30 minutes (min) to pull the enzyme solution into the leaf fragments, followed by shaking (30 rpm) for 2-3 hours then 60 rpm for 20 min at room temperature. The enzyme solution was pipetted out slowly of the petri dish into a 50 mL tube through a Falcon® 70 µm Cell Strainer with a 10 mL serological pipette to avoid breaking the cells. All centrifugation steps took place at room temperature, but tubes were kept on ice during all other steps. The protoplasts were spun down at 100 g for 5 min, then resuspended in 8 mL of W5 buffer (1.54 mL 5 M NaCl, 6.25 mL 1M CaCl_2,_ 125 μL 2M KCl, and 0.5 mL 0.2M MES), then layered on top of 5 mL of 0.55 M sucrose in a 15 mL tube, next centrifuged at 500 g for 5 min. The dark green layer above the sucrose and below the debris was removed and pipetted into a sterile 50 mL tube containing 10 mL of W5 buffer. This solution was centrifuged at 100 g for 5 min, and the cells were resuspended in 5 mL of W5. A small volume of the protoplast solution was used for imaging (check quality) and cell counting (for quantity). The cells were then spun down at 100 g for 5 min. MMG buffer (20 mL, 50 mL 0.8M <u>m</u>annitol, 2 mL 0.2 M <u>M</u>ES, 1.5 mL 1 M M<u>g</u>Cl_2,_ and 46.5 mL water) was added to the pelleted cells to have a final concentration of 1×10^6^ protoplasts per mL.

To transform protoplasts, 2.2 mL of protoplast solutions isolated above was added into a 50 mL tube containing 110 µg of plasmids of interest, mixed gently by tapping and then incubated in dark for 5 min. An equal volume of 40% PEG solution (4 g PEG 4000, 2.5 mL 0.8 M mannitol, 1 mL 1 M CaCl_2,_ and 3 mL water) was added to the tube and mixed by gentle inversion, followed by dark incubation on shaker (20-30 rpm) for 20 min. The W5 buffer (8.8 mL) was added to the tube to stop the transformation, followed by 5 min centrifugation at 100 g. This step was repeated two more times and the cells were resuspended in 2.2 mL of W5. Finally, 110 μL of Fetal Bovine Serum (FBS, Sigma F4135) and 11 μL of 10 mg mL^-1^ carbenicillin were added to the cells, which were kept in the dark overnight and transferred to low light the next morning.

Chlorophyll fluorescence in transformed protoplasts were measured at 25°C using a multi-wavelength kinetic spectrophotometer/fluorometer with a stirring enabled cuvette holder (standard 1 cm pathlength) designed and assembled by the laboratory of Dr. David Kramer at Michigan State University using the method described for algal cells with some modifications (Lucker & Kramer, 2013; Zhang *et al*., 2022). A 2.2 mL volume (around 12∼13 μg chlorophyll) of Setaria protoplasts were supplemented with 25 μL of fresh 0.5 M NaHCO_3_, loaded into a fluorimeter cuvette (C0918, Sigma-Aldrich), and dark-adapted for 10 min. Fluorescence measurements were taken with measuring pulses of 100 μs duration. The pulsed measuring beam was provided by a 505 nm peak emission light emitting diode (LED) filtered through a BG18 (Edmund Optics) color glass filter. The maximum efficiency of PSII (Fv/Fm) was measured with the application of a saturating pulse of actinic light with peak emission of 625 nm at the end of the dark adaptation period. After dark-adaptation, the protoplasts sample was illuminated by a pair of LEDs (Luxeon III LXHL-PD09, Philips) with maximal emission at 620 nm, directed toward both sides of the cuvette, perpendicular to the measuring beam. We conducted two kinds of measurements separately: (1) light responses curves from dark to 15, 35, 50, 100, 150 μmol photons m^−2^ s^−1^ light, each light lasted 3 min; (2) following the light response curve as in method 1, protoplasts samples were stay in 150 μmol photons m^−2^ s^−1^ for 15 min before 15 min dark, this methods allowed us to monitor NPQ induction in light and relaxation in dark. Isolated Setaria protoplasts were sensitive to light, thus, the maximum light of 150 μmol photons m^−2^ s^−1^ was used. NPQ was calculated as (F_m_-F_m’_)/F_m’_; PSII efficiency was calculated as (F_m_-F_o_)/F_m_ or (F_m’_-F_s_)/F_m’_ in dark or light adapted protoplasts, respectively (Baker *et al*., 2007; Zhang *et al*., 2022). F_m_ and F_m’_ were the maximum chlorophyll fluorescence in dark and light adapted protoplasts, respectively. F_o_ and F_s_ were the minimum and steady chlorophyll fluorescence in dark and light adapted protoplasts, respectively.

### Protein structure prediction

The structure predictions for the Arabidopsis and Setaria NPQ proteins are generated by MULTICOM3 (Liu *et al*., 2023a,b), which was built on top of AlphaFold2/AlphaFold-Multimer v2.2.0 (Jumper *et al*., 2021; Evans *et al*., 2022). Compared to the standard version of AlphaFold2/AlphaFold-Multimer v2.2.0, MULTICOM3 can improve the accuracy of tertiary structure prediction by 8-10% and quaternary structure prediction by 5-8% on the Critical Assessment of Structure Prediction (CASP15) dataset. For each protein, MULTICOM3 generated up to 55 structural predictions, from which the one with the highest AlphaFold2 pLDDT score was selected for display in Figure S2. The transmembrane domain predictions for the Arabidopsis and Setaria PsbS proteins are generated by DeepTMHMM v1.0.24 (Hallgren *et al*., 2022), a deep learning tool that can predict the topology of transmembrane proteins. For the dimer structure prediction of AT1G44575 (AtPsbS) and Sevir.5G400800 (SvPsbS), a total of 165 structure predictions were generated. The structure prediction with the highest AlphaFold-Multimer confidence score (0.7003) (Bryant *et al*., 2022) indicates a potential interaction between two proteins. See more information in Supplemental File 1.

### Generating transgenic Setaria lines

*Setaria viridis* WT (ME034) was used for all transformations. The Setaria AtVPZ lines were generated by the DARPA LISTENS team, with PvUbi2 promoter and AtHSP terminator for each of the three AtNPQ genes (*AtPsbS*, *AtVDE*, *AtZEP*) in one plasmid (Stone *et al*., 2024). The coding sequence of each Arabidopsis gene was codon optimized for the Setaria genome. The single-gene overexpression lines were generated by the Zhang Lab, with PvUbi2 promoter and AtHSP terminator for AtPsbS-overexpression (OE), AtVDE-OE, and AtZEP-OE lines, and pZmUbi1 and AtHSP terminator for SvPsbS-OE lines using the Golden Gate Cloning approach (Marillonnet & Grützner, 2020; Bird *et al*., 2022). Agrobacterium-mediated transformation in Setaria tissue culture was performed as described (Finley *et al*., 2021). All transgenic lines were selected using hygromycin and confirmed by genotyping.

### DNA isolation and genotyping

Two 2-cm segments of healthy leaf tissues were used for DNA extraction using a similar protocol as described (Chen & Ronald, 1999). Tissue was collected in a 1.75 mL tube with a grinding bead and frozen at –80°C until use. The tissue was homogenized using a TissueLyser II (QIAGEN, # 20.747.0001), followed by addition of 250 μL cetyltrimethylammonium bromide (CTAB) extraction buffer, incubated at 60°C for 30 min, and centrifuged at 10,000 g for 10 min (same speed for the rest of the centrifugation steps). The supernatant was transferred to a new tube with 2.5 μL of RNase A (10 mg mL^-1^), incubated at room temperature for 15 min, and then centrifuged for 5 min. The supernatant was added an equal volume of 24:1 chloroform/isoamyl alcohol, vortexed, and centrifuged for 1 min. The aqueous phase was transferred to a new tube with 0.7 volumes of cold isopropanol, incubated at –20°C for 20-30 min, and then centrifuged for 10 min. The supernatant was decanted, and the DNA pellet was washed with 70% ethanol then left to dry and finally resuspended in 20 μL of nuclease free water. DNA concentrations and quality were measured using dsDNA High-Sensitivity (HS) Qubit (Thermo Fisher Scientific Inc., #Q32854) and a NanoDrop 2000 Spectrophotometer (Thermo Fisher Scientific Inc., ND2000USCAN), respectively. Genotyping was performed using the TaqMan Genotyping Master Mix (Cat# 4371355, Life Technologies), reference probe (SiLeafy), target probe (hygromycin), reference primer set (ORZ937/938), and target primer set (ORZ875/876) (See primer sequences in Supplemental Table S1b). Leaf DNA of 15 ng in 8 μL were mixed with 12.4 μL of master mix for two technical replicates, thus 10 μL for each reaction in a Hard-Shell(R) 384-Well PCR Plate (Bio-Rad, # HSP3805). Three biological replicates were used for each genotype. Genotyped transgenic lines with confirmed zero, one (calibrator), two, or four copies of hygromycin genes were included as positive controls; water and WT DNA were used as negative controls. The qPCR reaction was performed in a CFX96 Touch Real-Time PCR Detection System (Bio-Rad) with the following protocol: 10 min at 95°C; then 50 cycles of 15 seconds at 95°C, 1 min at 60°C followed by a fluorescence reading. The cycle concluded with a cool down to 14°C for 5 min. The dual-channel probe option was used, detecting fluorophores HEX (SiLeafy) and FAM (hygromycin). HEX and FAM Cq values were used to calculate the number of hygromycin copies for each DNA sample. ⊗Cq was first calculated by subtracting the HEX Cq value from the FAM Cq value of the same sample. ⊗⊗Cq was calculated by subtracting the mean ⊗Cq of the single-copy calibrator technical replicates from the ⊗Cq value of the target sample. The number of hygromycin copies was calculated as 2^(-⊗⊗Cq)^.

### RNA isolation, cDNA synthesis, and quantitative real-time PCR (RT-qPCR)

Leaf tissue was collected from the middle 2-cm section of a fourth leaf of 14-day old plants. One leaf segment was collected from three different plants for each genotype into a 2 mL tube with a grinding bead (4039GM-S050, Inframat® Advanced Materials LLC), flash frozen in liquid nitrogen immediately after collection, and stored at –80°C until use. RNA was extracted as described before (Anderson *et al*., 2021). Leaf tissue was homogenized in a TissueLyser II (QIAGEN, # 20.747.0001), added 1 mL of TRIZOL (Thermo fisher, #15596026), and mixed well before 200 μL of 24:1 chloroform/isoamyl alcohol (Sigma-Aldrich, #C0549-1PT). The mixture was centrifuged for 15 min at 4°C and 11,000 rpm (the same speed and condition for all centrifugations). The top supernatant was transferred to a new tube, added equal volume of 24:1 chloroform/isoamyl alcohol, centrifuged for 5 min. The top supernatant was transferred to a new tube, added 0.7 volumes of cold 100% isopropanol, incubated at –20°C for 30 min, followed by 15 min centrifugation. Cold 75% ethanol was added to the pellet before centrifuging again for 2 min. The ethanol was decanted, and the last step was repeated. The pellet was left to dry, then resuspended in 50 μL of nuclease-free UltraPure water (Life Technologies, #10977015). RNA concentrations and quality were measured using Qubit RNA Broad Range (BR) Assay Kit (Thermo Fisher Scientific Inc., #Q10210) and a NanoDrop 2000 Spectrophotometer (Thermo Fisher Scientific Inc., ND2000USCAN), respectively. A 0.5 μg of RNA in 8 μL was used for cDNA synthesis using a SUPERSCRIPT III 1ST STRAND Kit (Thermo Fisher, #18080051) and cDNA was diluted to 1:20 for a 10-μL RT-qPCR reaction per well using the SensiFAST SYBR No-ROS kit (Bioline, BIO-98020). Two reference genes (UBIQ4 and BIND) were chosen based on their consistent expression in WT (Martins *et al*., 2016; Anderson *et al*., 2021). A primer set was designed to amplify each gene of interest: *AtZEP*, *AtVDE*, *AtPsbS*, *SvZEP*, *SvVDE*, and *SvPsbS* (See primer sequences in Supplemental Table S1b). Three biological replicates were used for each genotype, and three technical replicates were used for each biological replicate performed in a CFX384 Real-Time System (C 1000 Touch Thermal Cycler, Bio-Rad, Hercules, California). The RT-qPCR protocol was set up as follows: (1) 2 min at 95°C; (2) 40 cycles of 5 s at 95°C, 10 s at 60°C and 15 s at 72°C; (3) final melt curve, 5 s at 95°C, 5 s at 60°C, followed by continuous ramping of temperature to 99°C at a rate of 0.5°C s^−1^; (4) Cool down 30 s at 37°C. Melting curves and RT-qPCR products were checked to ensure there are no primer dimers or nonspecific PCR products. All qPCR products were sequenced to verify their identities by Eton Bioscience InC. The Cq value from two reference genes (UBIQ4 and BIND) was averaged for each sample. The mean Cq reference value was subtracted from the Cq value of the target gene for the same sample to calculate ΔCq. Relative expression was calculated by 2^(-ΔCq)^.

### Protein isolation and western blot

Two 2-cm segments from the middle section of a top fully expanded 4th leaf were collected from 14-day old plants for protein isolation. Leaf tissue was immediately frozen in liquid nitrogen after collection and stored at –80°C before use. Leaf tissue was homogenized using a grinding bead and TissueLyser II (QIAGEN, # 20.747.0001). Protein extraction buffer (1.25 mL 80% glycerol, 1.25 mL 0.5 M Tris-HCl, 2 mL 10% SDS, 0.5 mL 100% 2-mercaptoethanol, and 5mL water) was added to each sample equal to 50 mg fresh weight per mL. Samples were centrifuged at 13,000 g for 1 min at 4°C and the supernatant was transferred to a new tube. Protein concentrations were checked using a Pierce 660 nm kit. For western, a lower polyacrylamide SDS gel was made by mixing 2.5 mL of 4X lower buffer (1.5 M Tris pH 8.8 and 0.4% SDS), 2 mL 40% acrylamide, 5.5 mL sterile water, 100 μL 100% APS, and 10 μL TEMED. An upper 8% acrylamide SDS gel was made by mixing 4 mL 4X upper buffer (0.5 M Tris pH 6.8 and 0.4% SDS), 1 mL 40% acrylamide, 44 μL 10% APS, and 4.4 μL TEMED. Gel electrophoresis was performed, and the gel was transferred to a nitrocellulose membrane, both of which were surrounded on each side by five layers of Whatman filter paper. The gel, membrane, and filter paper were soaked in a 1X transfer buffer before assembling into layers. After the transfer, ponceau S stain was poured over the membrane and left for 5 min, then rinsed with water. Blocking was then performed by covering the membrane in milk solution (1 g milk powder, 20 mL 1X PBS buffer, and 0.1% tween) overnight at 4°C on a shaker. The 20X PBS buffer was made with 160 g NaCl, 4 g KCl, and 2.88 g Na_2_HPO_4_. After sitting overnight, the membrane was rinsed for 15 s two times with 10 mL of 1 x PBST (diluted from 20X PBS and 0.1% of tween). The primary antibody was diluted in PBST, poured onto the blot, and left to incubate for one hour at room temperature on a shaker. The membrane was then washed three times for 5 min each with PBST. The membrane was then treated with the secondary antibody (Anti-Rabbit IgG, Sigma, A9169, 1:10,000 dilution) for one hour at room temperature on a shaker and washed three times for 5 min each with PBST. To image the membrane, PICO solution was made by mixing 350 μL of each reagent and pouring over the membrane. A Chemiluminescence machine was used to image the western blots. The primary antibody used are all from Agrisera (Sweden): AtPsbS (AS09533, 1:2000 dilution), AtVDE (AS153091, 1:2000 dilution), AtZEP (AS153092, 1:500 dilution). We used 9 μg proteins per lane for westerns using antibodies of AtPsbS and AtVDE, but 27 μg proteins per lane for westerns using antibodies of AtZEP due to its low sensitivity. Western bands of interest were quantified from densitometry of the western blots using ImageLab Software, normalized to WT.

### Xanthophyll pigment analysis

Setaria plants grown under the control conditions for 14 days were subjected to fluctuating light treatment and the 4th leaves were harvested at one of three time points: (1) after 25 min dark adaptation in a dark chamber; (2) after time point 1 and 3 min at 1500 μmol photons m^−2^ s^−1^ inside of the LI-6800 leaf chamber; (3) after time point 1, and fluctuating light (1500, 200, 1500, 200, and 1500 μmol photons m^−2^ s^−1^, each light 3 min) followed by 5 min dark inside of the LI-6800 leaf chamber. Leaf segments inside the LI-6800 leaf chamber were collected in screw cap tubes (USA Scientific, #1420-9700), flash frozen in liquid nitrogen, and stored at –80°C until used. Xanthophyll pigments were quantified as described (Anderson *et al*., 2021). Three biological replicates of each genotype for each time point were collected. For pigment extraction, 600 µL of cold acetone was added to samples, then they were homogenized in a FastPrep-24 5G (MP Biomedicals, #116005500) at 6.5 m s^−1^ for 30 seconds at room temperature. Samples were centrifuged at 21,000 g for 1 min to remove cell debris. The supernatant was filtered through a 4 mm nylon glass syringe with a pore size of 0.45 µm (Thermo Scientific, #44504-NN). After filtration, samples were analyzed by HPLC on an Agilent 1100 separation module equipped with a G1315B diode array and a G1231A fluorescence detector. The data were collected and analyzed using the Agilent LC Open Lab ChemStation software. Pigments were separated on a ProntoSIL 200-5 C30, 5.0 μm, 250 mm × 4.6 mm column equipped with a ProntoSIL 200-5-C30, 5.0 μm, 20 mm × 4.0 mm guard column (Bischoff Analysentechnik). De-epoxidation state was calculated by (zeaxanthin + 0.5 antheraxanthin) / (violaxanthin + antheraxanthin + zeaxanthin), assuming interconversion of the intermediate antheraxanthin between zeaxanthin and violaxanthin.

### Gas exchange and chlorophyll fluorescence measurements

A gas-exchange system LI-6800 with a Fluorometer head 6800-01 A was used to measure pulsed amplitude modulated chlorophyll a fluorescence (LI-COR Biosciences, Lincoln, NE) as described with some modifications (Anderson *et al*., 2021). The environmental parameters in the LI-COR leaf chamber were maintained at 400 ppm CO_2_, 25°C leaf temperature, 500 μmol s^−1^ flow rate, 1.5 kPa leaf VPD and 10,000 RPM fan speed for all measurements. Before LI-COR measurements, a 14-day old Setaria plant was dark-adapted in a dark chamber inside of a growth chamber with the control conditions for 20 min. Then a dark-adapted, 4^th^ intact leaf was put into the LI-6800 chamber for an extra 5 min in dark, followed by a pulse of saturating light to measure maximum PSII operating efficiency (F_v_/F_m_). F_m_ and F_v_ are the maximum and variable chlorophyll fluorescence in dark-adapted leaves (Maxwell & Johnson, 2000; Baker *et al*., 2007). For the fluctuating light protocol, plants were exposed to alternating cycles of 1500, 200, 1500, 200, 1500 μmol photons m^−2^ s^−1^ light. Each light phase lasted 3 min, with gas exchange and chlorophyll fluorescence measured every 1 min. PSII operating efficiency was measured as (1-F_s_/F_m_’); F_s_ and F_m_’ are the steady and maximum chlorophyll fluorescence in light-adapted leaves. The fluctuating light protocol was followed by 5 min darkness to measure NPQ relaxation, with chlorophyll fluorescence measured every 30 seconds. The NPQ dark relaxation was performed in several batches of plants so the chlorophyll fluorescence measurements were shifted by 5 s relative to the light-off time point and with 5 s time shift interval between two sequential batches as described (Kromdijk *et al*., 2016). Each batch has at least 3 biological replicates. NPQ in light and post-light dark was calculated as (F_m_/F_m_’-1) and (F_m_/F_m_’’-1); F_m_’ and F_m_’’ are the maximum chlorophyll fluorescence in light-adapted leaves and post-light leaves in dark, respectively (Maxwell & Johnson, 2000; Baker *et al*., 2007). NPQ values during post-light dark were normalized to NPQ values just before dark relaxation within each set. Normalized NPQ values of all batches for each genotype were compiled as a function of time to generate time-series with a 5 s resolution, which was fitted by 1^st^ or 2^nd^ order exponential decay using the OriginLab software to get the decay time constants. For the light response curves, a dark-adapted leaf was exposed to increasing light intensities, 100, 200, 400, 600, 800, 1200, 1500 μmol photons m^−2^ s^−1^, with 3 min for each light intensity and one measurement per min.

### MultispeQ measurement

Photosynthetic parameters were measured using a MultispeQ v2.0 (Kuhlgert *et al*., 10/2016; Anderson *et al*., 2021) (PhotosynQ, East Lansing, MI, USA) using intact, light-adapted Setaria leaves inside the control growth chamber. The MultispeQ was modified with a light guide mask to improve measurements on smaller leaves. Measurements were taken within 15 s at room temperature. Electrochromic shift (ECS), a useful tool for measuring proton fluxes and the transthylakoid proton motive force (*pmf*) *in vivo*, was measured through light to dark transition induced electric field effects on carotenoid absorbance bands (Witt, 1979; Baker *et al*., 2007). The decay time constant (−_ECS_) of light–dark-transition-induced ECS signal is inversely proportional to proton conductivity (ɡ_H_^+^ = 1/−_ECS_), which is proportional to the aggregate permeability of the thylakoid membrane to protons and largely dependent on the activity of ATP synthase. The proton flux rate (v_H+_) was calculated by ECS_t_/−_ECS_. Photosynthetic parameters were measured at 250, 500, and 1000 μmol photons m^−2^ s^−1^ using a modified photosynthesis RIDES protocol.

### Plant stress treatments

Setaria plants were grown under control conditions for nine days with normal watering as mentioned above. Then some plants stayed in the control condition while others were subjected to different stress treatments for five days, including high temperature (constant 40°C), drought (no watering), high light (950 µmol photons m^−2^ s^−1^), low light (100 µmol photons m^−2^ s^−1^), and greenhouse conditions (dynamic changes of environmental conditions, August 2022 and July 2024, St. Louis, USA). For stress treatments in growth chambers, other unmentioned environmental parameters stayed the same as the control condition. When plants were 14 days old under either the control or stressful conditions, images were taken of each plant, and wet and dry biomass of the entire plant (from base just above the soil) were quantified. Dry biomasses were measured by placing the whole above-ground plant in a drying oven at 60°C for six to seven days. Plant height, from plant base to the highest stem, was quantified using plant images using ImageJ. Several rounds of stress treatment were performed. Within one round of treatment, each plant parameter was normalized to the mean values of WT plants grown under the same conditions.

### Chlorophyll extraction

The 4^th^ fully expanded leaf was collected from 14-day old plants. The fresh weight of leaves was measured. Leaves were harvested in a tube containing a grinding bead and put into liquid nitrogen, then stored at –80°C. For chlorophyll extraction, samples were grinded in a TissueLyser II (QIAGEN, # 20.747.0001). One mL of 80% acetone was added to the homogenized tissue and mixed thoroughly. Samples were centrifuged at 3,000 g for 5 min at 4°C. The supernatant (∼200 μL) was added to 800 μL of 80% acetone. This mixture was added to a cuvette and measured using a spectrophotometer. One mL of 80% acetone was used as a blank and wavelengths of 663 nm (chlorophyll a), 646 nm (chlorophyll b), and 470 nm (carotenoids) were measured. Chlorophyll content was calculated using the following formula (Wellburn, 1994): [(12.21xA663) – (2.81xA646)] x DF/FW where DF is dilution factor (which is 5) and FW is fresh weight. Chlorophyll b content was calculated using the following formula: [(20.13xA646) – (5.03 x A663)] x DF/FW. Carotenoid content was calculated using the following formula: [(1000 x A470) – (3.27 x Chl a) – (104 x Chl b)/198] x DF/FW. All calculated pigment units are μg mL^-1^.

### ABA quantification

After control and stress treatments, a 4th leaf was harvested for ABA measurement as described before (McAdam *et al*., 03/2016; Anderson *et al*., 2021). Frozen samples were homogenized, and 15 ng of [^2^H_6_]-abscisic acid was added as an internal standard. Samples were placed under a vacuum to be dried completely, resuspended in 200 µL of 2% acetic acid in water (v/v), centrifuged, and an aliquot was used for ABA quantification. Measurements of foliar ABA levels were quantified using liquid chromatography tandem mass spectrometry with an added internal standard via Agilent 6400 Series Triple Quadrupole liquid chromatograph associated with a tandem mass spectrometer.

### Statistical analysis

For most cases, we used a two tailed *t*-test assuming unequal variance for statistical analysis, with a significance level as of P<0.05. To compare light response curves with lots of data points as in Figure S8, S9, we analyzed the whole curves but not individual data points by performing statistical modeling and posterior probabilities. For long LICOR experiments with gradually increased light intensity (Fig. S8), the data of NPQ was modeled using a monomolecular growth curve with an intercept term; the data of PSII operating efficiency was modeled as monomolecular decay from an initial starting value using Student’s *t*-distribution for robust regression; the data of net CO_2_ assimilation rate was modeled using a Student’s *t*-distribution as monomolecular growth with a three parameter logistic submodel for the sigma parameter. MultiSpeQ data were analyzed using Bayesian hierarchical growth (or decay) models (Fig. S9): ECS_t_ data was modeled as a linear trend; g_H+_ data was modeled as a power law growth curve with an intercept term; the data of v_H+_ and PS1 oxidized centers were modeled as a power law growth curve with an intercept term. All models were fit using 4 chains, with each chain running 1000 burn-in iterations and 1000 sampling iterations. Posterior distributions from the fit models were compared between parameters for all genotypes testing for a difference in posterior distributions with posterior probability of at least 95% as significance. All the analyses mentioned were run in R version 4.3.2 running on Ubuntu 22.04. We used the pcvr (version 0.2.0), brms (version 2.21.0), ggplot2 (version 3.5.1), and readxl (version 1.4.3), and cmdstanr (version 0.6.1) in R packages as well as CmdStan version 2.33.1 for model fitting (Wickham, 2009; Bürkner, 2017; Wickham H, 2023; Summer, 2024).

## Results

To investigate the function of Arabidopsis NPQ orthologs in Setaria, we first overexpressed codon-optimized Arabidopsis NPQ genes in Setaria protoplasts (Fig. 1). Freshly isolated Setaria protoplasts were transformed with cassettes containing Arabidopsis NPQ genes, then recovered in dark for 24 h before chlorophyll fluorescence measurements using a kinetic spectrophotometer/fluorometer. Due to the light sensitivity of isolated protoplasts, we performed light responses curves from dark to a maximum light of 150 μmol photons m^−2^ s^−1^. In response to increased light, *AtVDE*-OE protoplasts (with overexpressed *AtVDE*) induced a greater amount of NPQ at lower light intensities than the negative controls with water (Fig.1a). *AtZEP*-OE protoplasts had a slight reduction of NPQ as compared to the water control. Surprisingly, *AtPsbS*-OE protoplasts had significantly reduced NPQ. The differences in PSII operating efficiency were small among all constructs except for a significant decline in *AtPsbS*-OE protoplasts (Fig.1b). To understand temporal responses of NPQ induction and relaxation in transformed Setaria protoplasts, we extended the previous light response curves by adding a 15-min constant light at 150 μmol photons m^−2^ s^−1^, followed by 15 min in dark (Fig.1c, d). This long protocol gave similar results as the short protocol. And *AtZEP*-OE protoplasts had increased PSII efficiency than the water control. The results suggest AtVDE and AtZEP proteins may behave similarly in Setaria as in C_3_ plants; however, the AtPsbS protein may function or be regulated differently in C_4_ plants and overexpressing *AtPsbS* may be unfavorable in Setaria protoplasts.

**Figure 1.**
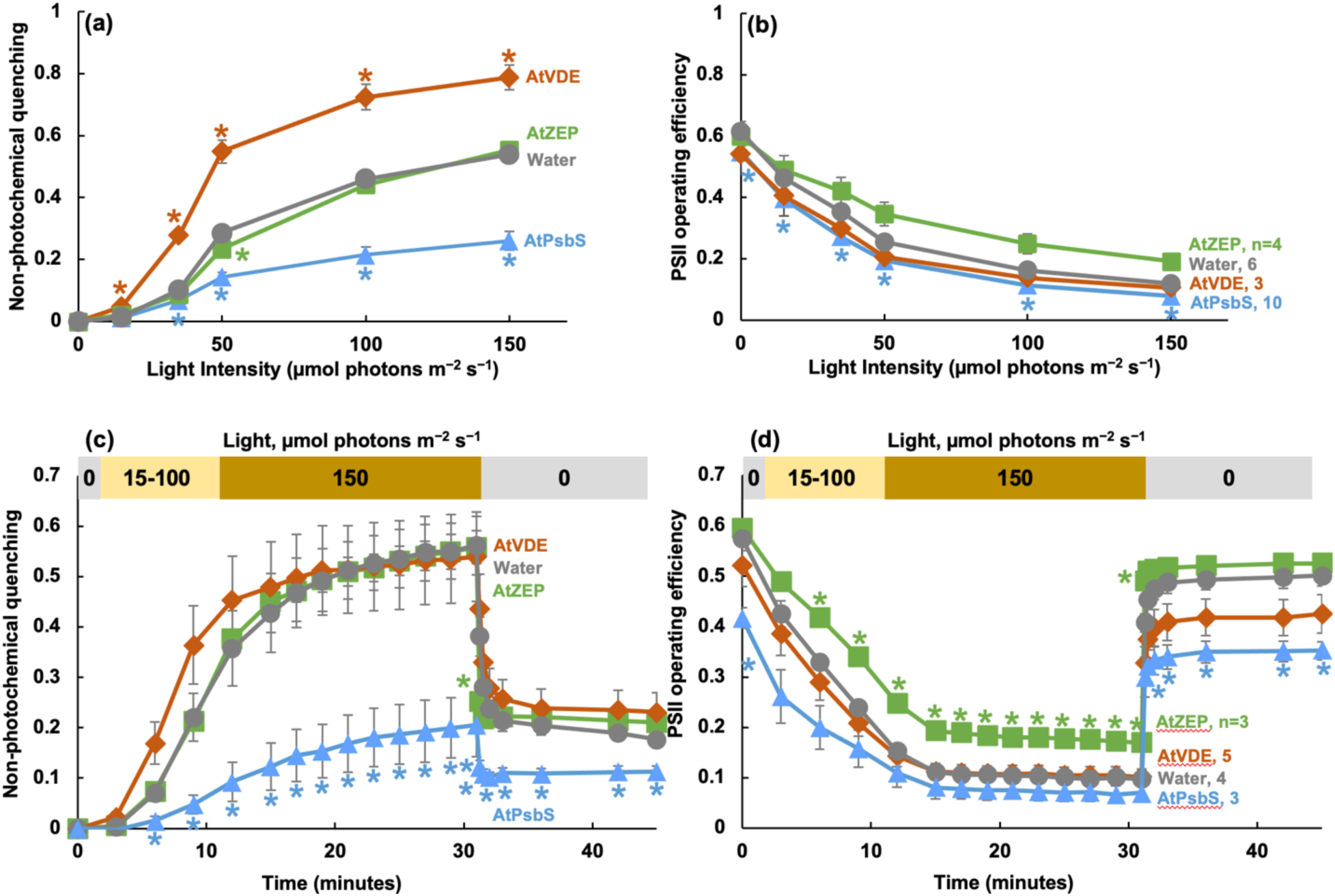
Overexpression of AtPsbS in Setaria protoplasts resulted in reduced NPQ and PSII operating efficiency. Isolated Setaria protoplasts were transformed with constructs overexpressing one of the indicated NPQ genes from Arabidopsis (*AtVDE*, *AtZEP*, *AtPsbS*). Protoplasts transformed with water served as negative controls. Chlorophyll fluorescence was measured in protoplasts 24 h after the transformation. **(a, b)** Transformed, dark-adapted protoplasts were subjected to a series of increased light intensity of 0, 15, 35, 50, 100, 150 µmol photons m^−2^ s^−1^, each light lasted 3 min. **(c, d)** Transformed, dark-adapted protoplasts were subjected to a series of increased light intensity of 0, 15, 35, 50,100 µmol photons m^−2^ s^−1^, each light lasted 3 min, then illuminated with 150 µmol photons m^−2^ s^−1^ light for 15 min, followed by 15 min darkness to monitor NPQ decay. *, P<0.05, compared to protoplasts transformed with water using a Student’s two-tailed *t*-test assuming unequal variance. Mean ± standard errors (SE), n=3-10. Some error bars are too small to see. The replication numbers were marked in panel **b** and **d**.

Protein alignments and structural predictions of Setaria and Arabidopsis NPQ proteins show high similarity in sequences and structures (Fig. S1, S2). Like the AtPsbS protein, SvPsbS protein also has four transmembrane domains and the two conserved glutamate residues for pH sensing (Fig. S1, S3) (Li *et al*., 2004). However, our computational structure prediction suggests that AtPsbS can heterodimerize with SvPsbS to form a dimer, and that their binding is tighter than any other PsbS pairs we tested, including the expected homodimers of AtPsbS or SvPsbS (Fig. 2). The Sv/At, At/At, Sv/Sv PsbS dimer pairs were predicted to have 11, 9, 7 interacting sites, respectively, with the pH-sensing glutamates either in or near the interacting sites (Fig. 2a, b, S3b, c, Supplemental file 1).

**Figure 2.**
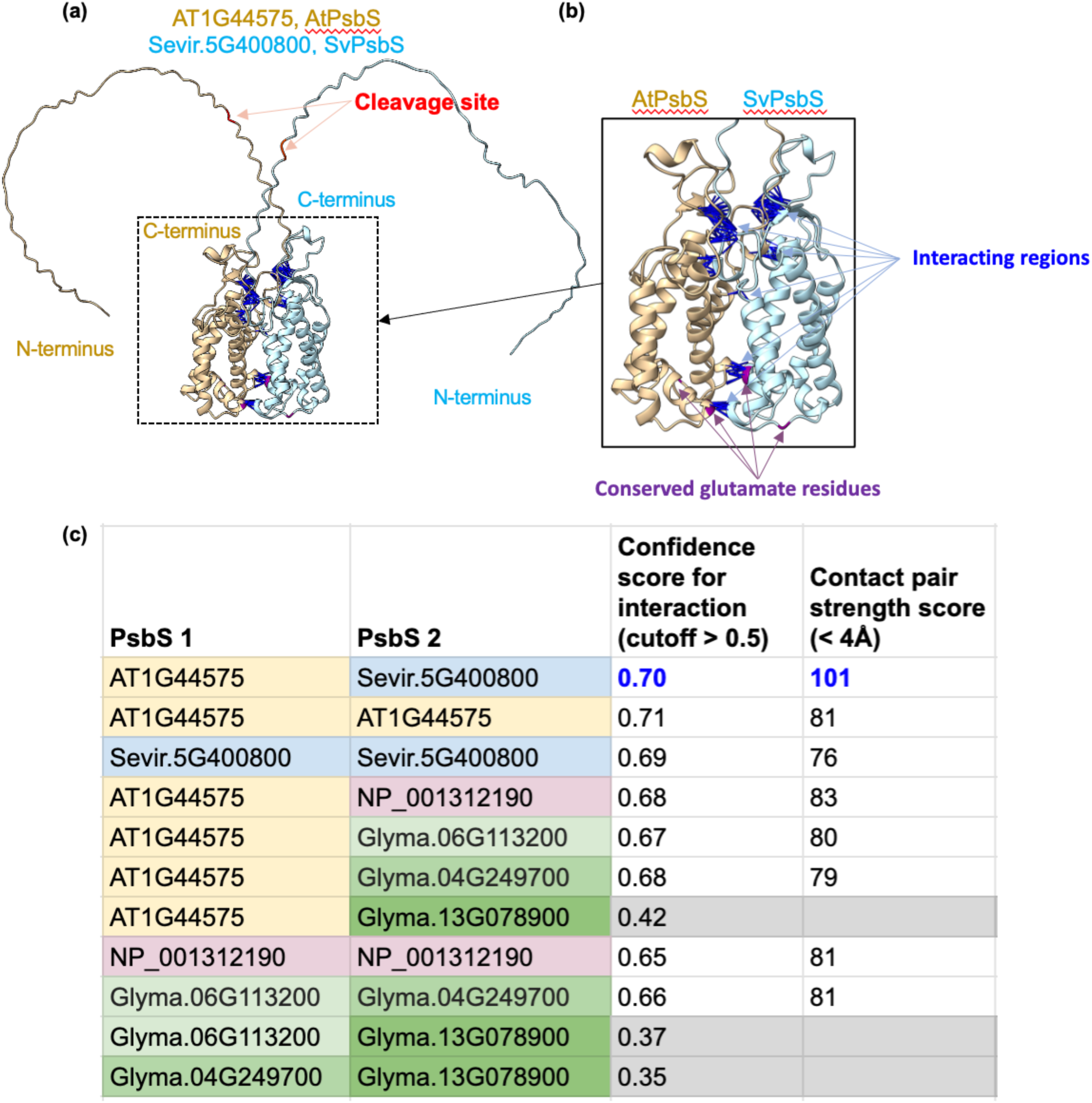
The dimer structure predictions of AtPsbS and SvPsbS suggest a potentially tight interaction. **(a)** AtPsbS (brown) and SvPsbS (cyan) are predicted to interact with each other. The predicted cleavage site for chloroplast transit peptide of each protein is highlighted in red. The N and C terminus of each protein are marked. The predicted interacting regions are marked by blue lines. Due to the visualization angle of the predicted structure, only some interaction clusters are shown. **(b)** A close view of AtPsbS (brown) and SvPsbS (cyan) interaction sites (blue). The conserved glutamate residues of PsbS that sense pH are marked in purple. **(c)** AtPsbS and SvPsbS were predicted to have tighter interactions than other PsbS pairs. The interaction confidence scores and interaction strength scores of protein pairs were predicted by MULTICOM3, a protein complex structure prediction system powered by AlphaFold-Multimer. The interaction confidence scores have a cutoff of >0.5, the bigger a score, the higher confidence for the predicted interaction, scores of <0.5 means no confident interaction. Contact pair strength score is the number of interacting residue pairs that have a minimum distance < 4Å (angstrom, 0.1 nm). The bigger a contact pair strength score, the tighter the predicted interaction between two proteins. At, Arabidopsis thaliana; Sevir, Setaria viridis; NP, Nicotiana tabacum, tobacco; Glyma, Glycine max, soybean. Arabidopsis, Setaria, tobacco all have one PsbS protein. Soybean has three copies of the PsbS protein.

To assess the physiological relevance of the observed protoplast phenotypes and computational prediction, we next generated stable transgenic lines overexpressing one of the three Arabidopsis NPQ genes (AtNPQ lines, called AtVDE, AtZEP, or AtPsbS lines) and employed AtVPZ lines that overexpressed all three NPQ genes (Stone *et al*., 2024) (Fig. 3a). The Setaria AtVPZ lines we used had minimal phenotyping characterization previously (Stone *et al*., 2024). Because of the surprisingly reduced NPQ results when overexpressing the *AtPsbS* gene in Setaria protoplasts (Fig. 1), we also generated stable transgenic lines overexpressing Setaria *PsbS*, called SvPsbS lines. Homozygous overexpression lines were identified using genotyping qPCR. Most lines had reasonable chances for identification of homozygous lines (11%-25%), except for AtPsbS: we only identified one homozygous AtPsbS line out of 96 T_1_ or T_2_ plants screened, which is around 1% (Table 1). The result suggests that overexpression of AtPsbS in Setaria may be costly to overall fitness.

**Figure 3.**
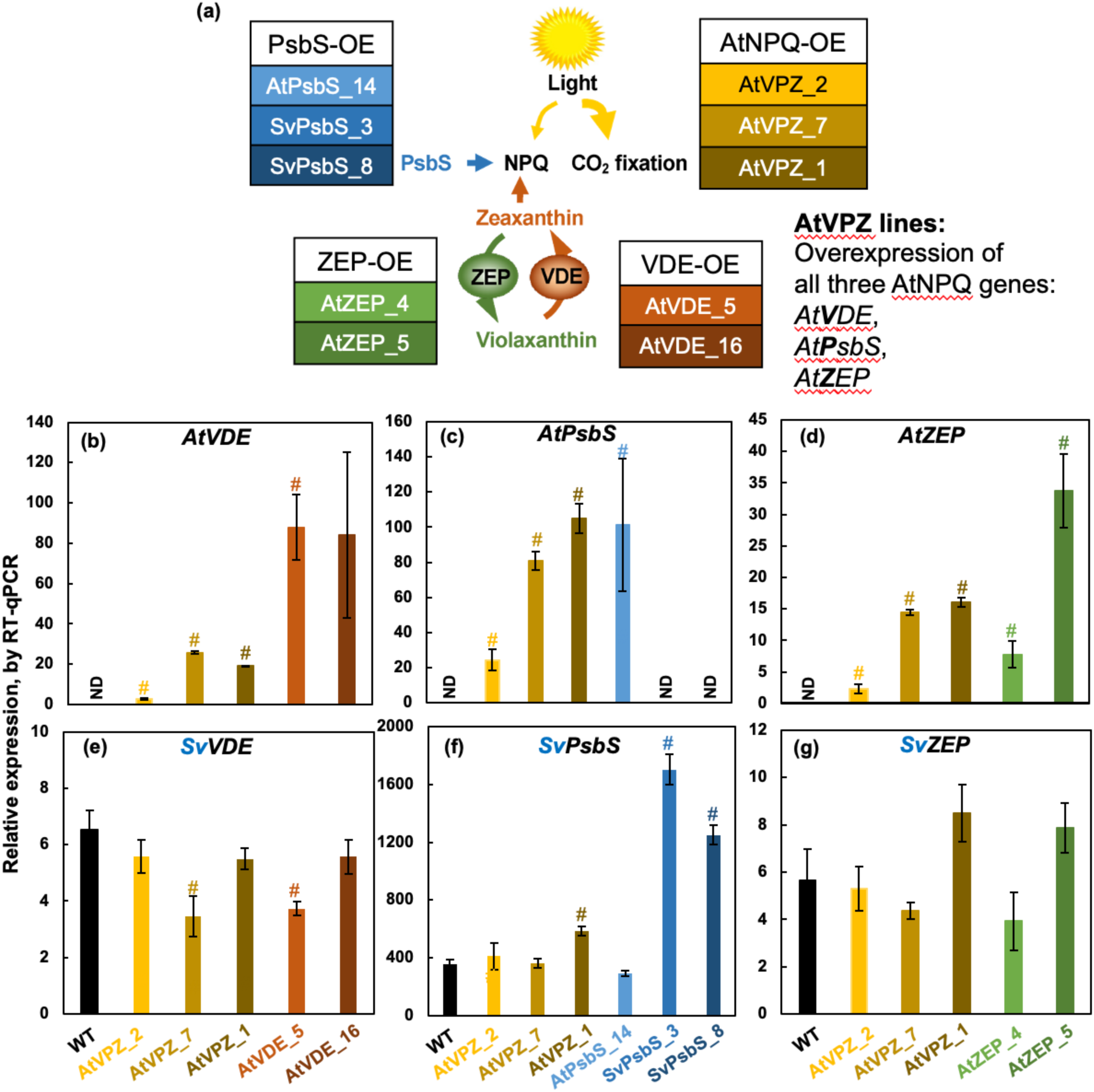
Setaria homozygous transgenic lines with overexpressed NPQ genes were confirmed at the transcript level through RT-qPCR. **(a)** Simplified NPQ pathways of NPQ and zeaxanthin formation as well as our related Setaria transgenic lines. Overexpression, OE. Each table contains names of transgenic lines that overexpress the indicated genes. At, genes from Arabidopsis. Sv, genes from Setaria. AtVPZ lines are Setaria transgenic plants that overexpress all three of the Arabidopsis NPQ genes: *At**V**DE*, *At**P**sbS*, and *At**Z**EP*. Independent transgenic lines are indicated by the numbers after a dash. The box color for transgenic lines in the tables match the color of lines and bars representing these transgenic lines in the rest of our figures. (**b-g**) Relative expression based on RT-qPCR results for the indicated transcript labeled at the top of each panel: Arabidopsis (**b, c, d**) and Setaria (**e, f, g**) NPQ genes, in Setaria WT and transgenic lines. *, P<0.05; #, P<0.01, transgenic lines were compared to WT for each indicated transcript using a Student’s two-tailed *t*-test assuming unequal variance. Mean ± SE, n=3. ND, not detected.

**Table 1.** Homozygous transgenic lines with AtPsbS overexpression were under-represented. We screened and genotyped T_1_ and T_2_ plants from T_0_ parent plants with one copy of hygromycin (hygro) gene using qPCR. We expect about 25% of plants from a parent plant with one hygro copy to be homozygous (with two copies of hygro gene).

| Overexpressed gene | Total T1 or T2 plants screened | # of plants with 0, 1, and 2 hygro copies | % of plants with 0 hygro copy | % of plants with 1 hygro copy | % of plants with 2 hygro copies |
| --- | --- | --- | --- | --- | --- |
| *SvPsbS* | 37 | 8, 21, 8 | 22% | 57% | 22% |
| *AtPsbS* | 96 | 78, 17, 1 | 81% | 18% | 1% |
| *AtVDE* | 22 | 11, 8, 3 | 50% | 36% | 14% |
| *AtZEP* | 35 | 16, 10, 9 | 46% | 29% | 26% |
| *AtVPZ* | 18 | 13, 3, 2 | 72% | 17% | 11% |

We then used RT-qPCR to check the RNA expression level of overexpressed genes in T_4_ or T_5_ homozygous Setaria lines (Fig. 3b-g). The three AtVPZ lines had increased transcript levels of all three NPQ genes compared to WT, although different lines varied in the induction level. The AtVPZ-2 line had the smallest magnitude of overexpression of the three NPQ genes, whereas AtVPZ-7 and AtVPZ-1 lines had higher, but similar expression levels between the two lines. *AtZEP*, *AtVDE*, *AtPsbS,* and *SvPsbS* transcripts had the expected increase in corresponding single trait overexpression lines as compared to WT. The AtNPQ lines with single gene overexpression often had much higher induction of the targeted transcripts than that in the AtVPZ lines. Additionally, the effects of overexpression on the native Setaria NPQ transcripts were mostly minimal, though we observed reduced SvVDE transcript levels in AtVPZ-7 and AtVDE-5 lines and increased SvPsbS transcripts in AtVPZ-1 lines (Fig. 3e-g).

We also quantified the abundances of overexpressed NPQ proteins in these homozygous transgenic Setaria lines using western blots (Fig. 4). The VDE antibody used showed specificity to the Arabidopsis NPQ orthologs, but the PsbS and ZEP antibodies may not distinguish the Arabidopsis and Setaria versions of these proteins. The three AtVPZ lines had increased but variable levels of all three NPQ proteins as compared to WT. The AtVPZ-2 line had the least induction of the three NPQ proteins and the AtVPZ-1 line had the highest induction of ZEP protein. The AtVDE, AtZEP, and SvPsbS lines with single gene overexpression often had much higher induction of the targeted proteins than that of the AtVPZ lines, except for the AtPsbS line. The induced NPQ protein levels were often consistent with the induced corresponding transcription levels (Fig. 3), except for the *AtPsbS* transcripts. Considering the significant induction of *AtPsbS* transcripts but much lower than expected of AtPsbS proteins (only 1.5 X folds as compared to WT) in these lines (Fig. 3c, 4), it may evidence post-transcriptional repression or degradation of the AtPsbS protein in Setaria.

**Figure 4.**
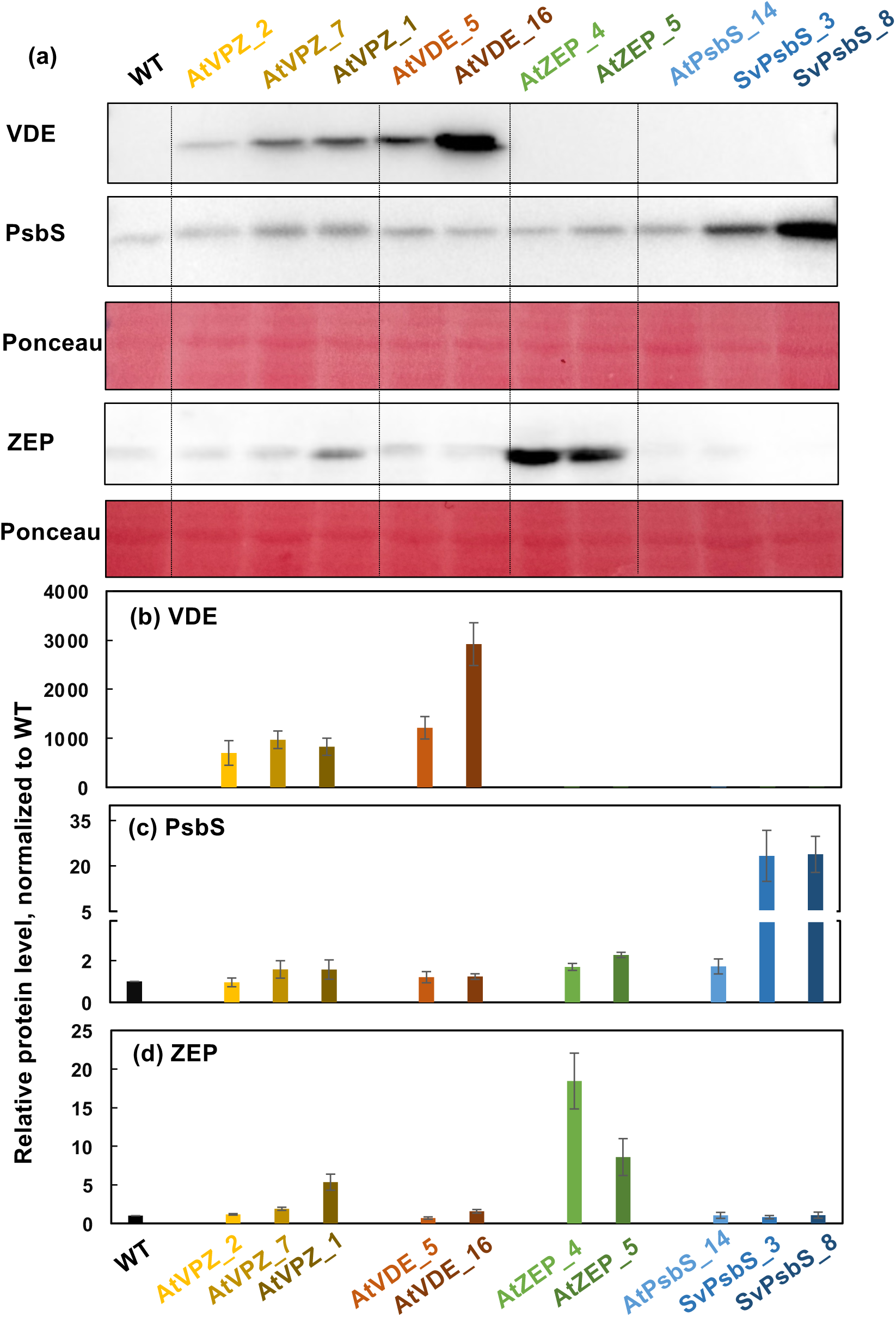
Setaria transgenic lines were confirmed at the protein level using western blots. The names for transgenic lines were the same as in Figure 3. **(a)** Representative western blot analysis of protein levels. The blotting for AtVDE and AtPsbS proteins were performed in the same run with all lines (9 µg protein per lane); the blotting for AtZEP protein was performed in a different run due to low sensitivity of the ZEP antibody which required increased protein loading (27 µg protein per lane). Ponceau stain was used as a loading control. **(b-d)** Protein level quantification relative to WT, determined from densitometry of western results. Mean ± SE, n=3.

Under the control growth conditions, these transgenic lines had little changes in chlorophyll or carotenoid contents as compared to WT (Fig. S4). In dark-adapted leaves, the maximum PSII efficiency of these transgenic lines was mostly similar to WT, except for AtVPZ-2 and SvPsbS-3 (higher and lower than WT, respectively) (Fig. S5a). AtVPZ-1 and AtVDE-16 lines had lower minimal chlorophyll fluorescence F_o_ than WT (Fig. S5b); while AtVPZ-2 and AtVPZ-7 had higher maximum chlorophyll fluorescence F_m_ and the AtVDE lines had lower F_m_ than WT (Fig. S5c). The lower F_o_ and F_m_ in the two AtVDE lines than WT may suggest quenching of chlorophyll fluorescence even in dark-adapted leaves of the AtVDE lines.

We further characterized these homozygous transgenic Setaria lines by xanthophyll pigment analysis (Fig. 5). Dark-adapted, intact leaves were subjected to fluctuating light between 200 (control light) and 1500 (high light) μmol photons m^−2^ s^−1^ in a LI-6800 leaf chamber (Fig. 5a). After 25 min dark acclimation and prior to actinic light treatment, the AtVDE lines still retained substantial amounts of zeaxanthin and antheraxanthin levels in contrast to all other lines (Fig. 5b, c), which may correlate with the lower F_o_ and F_m_ in the AtVDE lines as mentioned above (Fig. S5). With a 3-min high light treatment, zeaxanthin was quickly induced to above detectable level in most lines (except for AtZEP-4), with highest levels in AtVDE lines, followed by two AtVPZ lines (line 7 and 1). After 5-min dark acclimation following the fluctuating light regime (TP3), two of the AtVPZ lines (line 7 and 1) and two AtVDE lines had much higher zeaxanthin levels than WT, consistent with overexpressed *AtVDE* gene, while the two AtZEP lines had significantly reduced zeaxanthin levels as compared to WT, consistent with the accelerated zeaxanthin epoxidation due to the overexpression of *AtZEP*. Additionally, the AtVDE and AtVPZ lines had reduced violaxanthin levels as compared to WT after the 5-min post-light dark treatment (Fig. 5d), consistent with their increased zeaxanthin levels and overexpressed AtVDE proteins. The de-epoxidation states were consistent with zeaxanthin abundances (Fig. 5e). These results strongly support that: (1) the overexpressed *AtVDE* and *AtZEP* worked as expected *in vivo* in Setaria; (2) overexpression of *AtPsbS* or *SvPsbS* did not significantly affect zeaxanthin levels in Setaria; and (3) the AtVPZ and AtVDE lines had more zeaxanthin than WT even after 5-min dark recovery.

**Figure 5.**
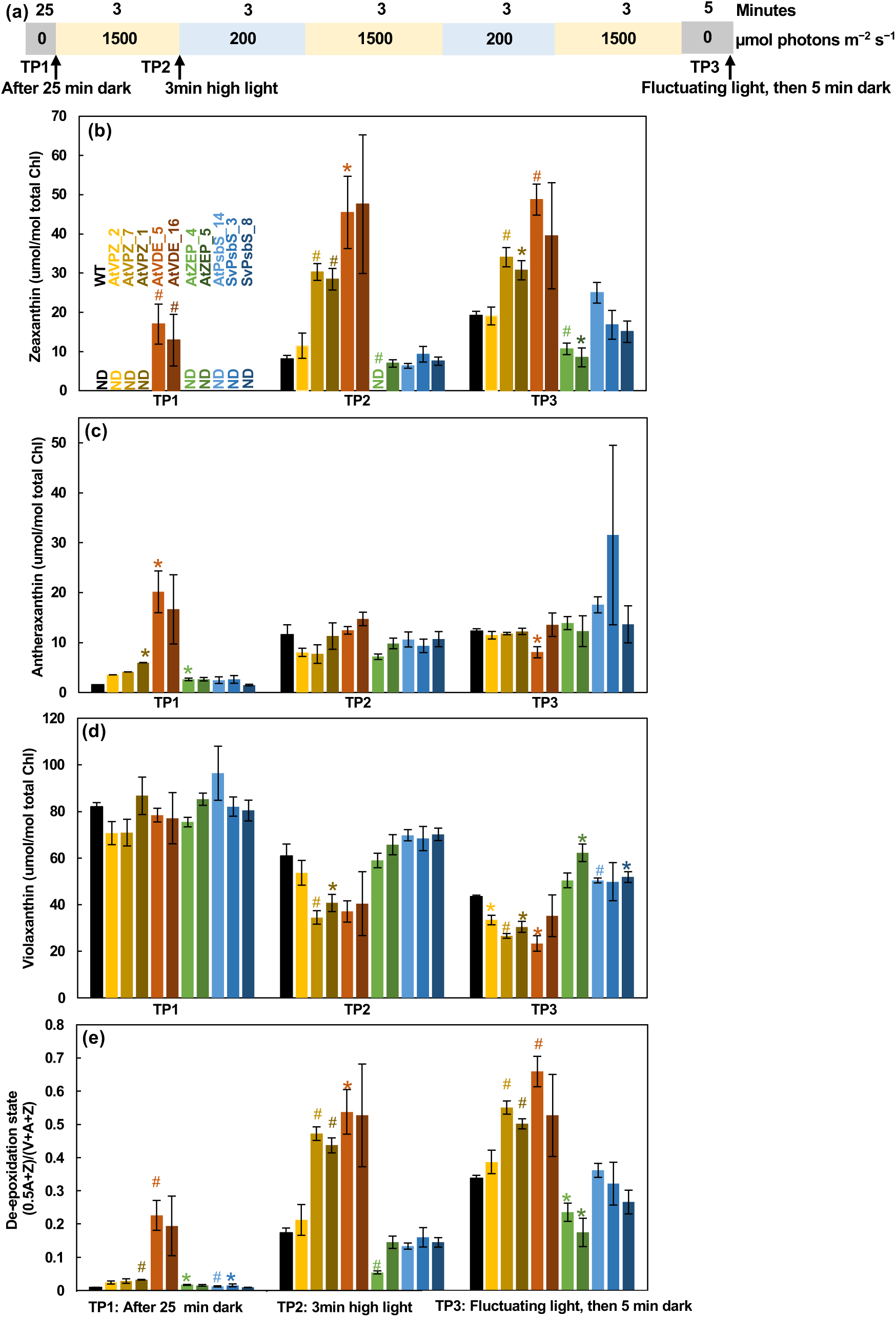
Setaria transgenic lines were confirmed at the pigment level using high-performance liquid chromatography (HPLC). **(a)** Leaves were collected after three time points (TP): 25 min in dark, 3 min at 1500 µmol photons m^−2^ s^−1^ light, and fluctuating light conditions followed by 5 min in dark. **(b, c, d, e)** The levels of zeaxanthin (Z), antheraxanthin (A), violaxanthin (V), and the xanthophyll cycle de-epoxidation state in *Setaria* WT and transgenic lines. *, P<0.05; #, P<0.01, transgenic lines were compared to WT under the same condition using a Student’s two-tailed *t*-test assuming unequal variance. Mean ± SE, n=3.

Overexpression of *AtVDE*, *AtZEP*, *SvPsbS*, but not *AtPsbS*, had expected effects on NPQ induction kinetics under the fluctuating light condition (Fig. 6). Two of the three AtVPZ lines (AtVPZ-1 and –7) had faster NPQ induction and higher NPQ levels than WT during the high-light phase of the fluctuating light experiment, whereas AtVPZ-2 had WT-like induction capacity but lower residual NPQ during the second low light phase (Fig. 6b). The AtVDE and SvPsbS lines also had faster NPQ induction and higher NPQ level than WT during the high-light phase of the fluctuating light experiment, while the two AtZEP lines had lower NPQ levels, as expected (Fig. 6c-e). However, the AtPsbS line had WT-level NPQ (Fig. 6e), which may correlate with the lower PsbS protein abundances observed (Fig. 4a). The magnitude of NPQ phenotypes in these transgenic lines were consistent with their induced zeaxanthin levels during the fluctuating light treatment (Fig 5). AtVPZ lines, and in particular AtVPZ_2 (p < 0.01), had higher net CO_2_ assimilation rates than WT under the fluctuating light condition, but no significant differences in PSII operating efficiencies across all these transgenic lines as compared to WT (Fig. S6).

**Figure 6.**
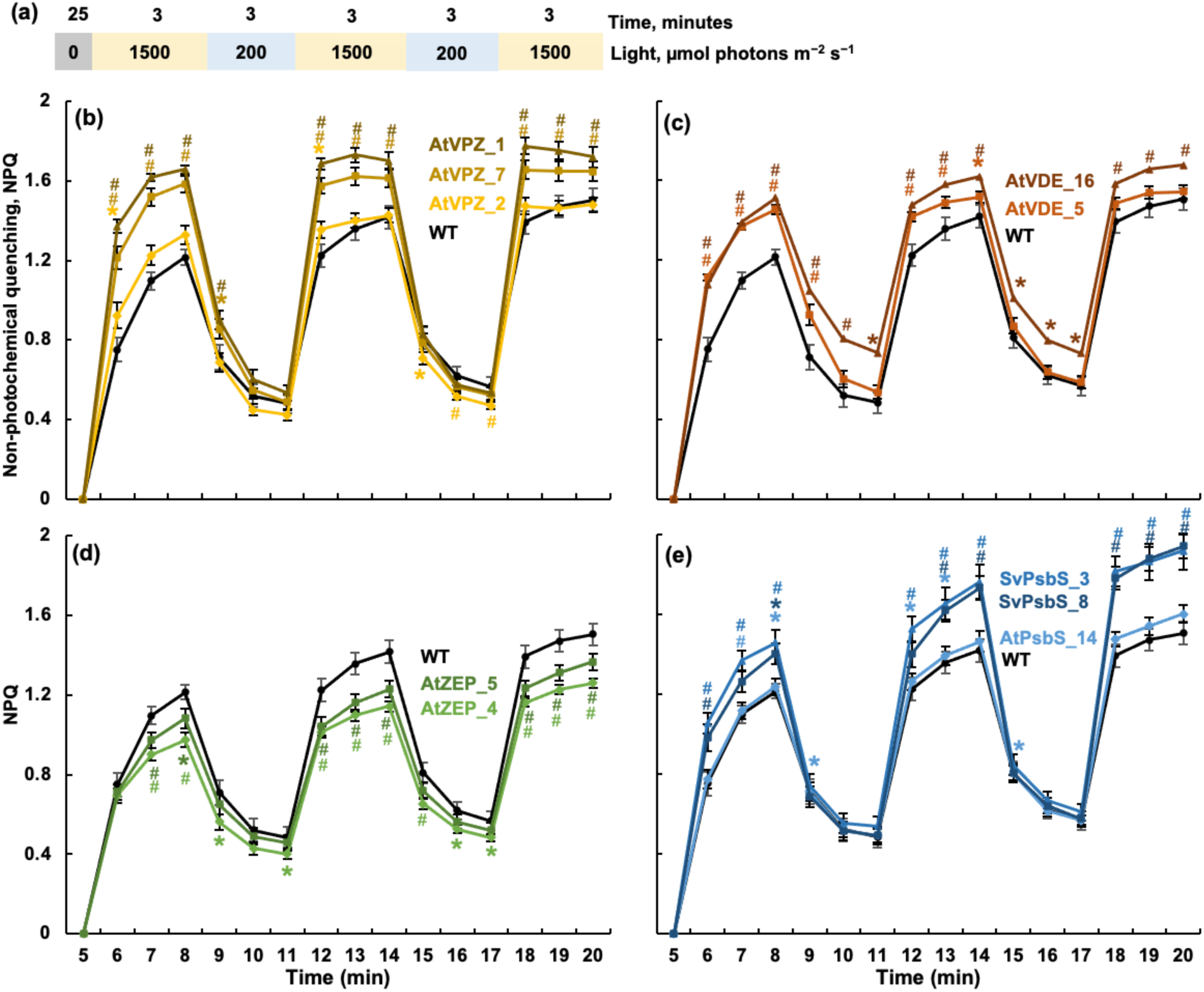
Setaria transgenic lines were confirmed through NPQ measurements. Plants were dark-adapted for 25 min, then a fully expanded 4th leaf was used for NPQ measurements by chlorophyll fluorescence in a LI-6800 using the fluctuating light treatment as shown in **(a)**. Each light phase lasted 3 min. **(b-e)** NPQ of WT and transgenic plants during fluctuating light conditions. Genotype names were labeled by the corresponding curves with the same color as the curve. *, P<0.05; #, P<0.01, transgenic lines were compared to WT under the same condition using a Student’s two-tailed *t*-test assuming unequal variance. Mean ± SE, n=18-23.

Overexpression of NPQ genes affected the normalized rate of NPQ relaxation during the 5-min dark treatment following the fluctuating light regime in Setaria (Fig. 7). The NPQ decay time constant (inverse to NPQ decay rate) was quantified using the 1^st^ order exponential decay. AtVPZ and AtVDE lines had increased NPQ decay time constants, which means slower NPQ decay in the dark (Fig. 7a, b, e), consistent with their higher zeaxanthin levels than WT after 5 min post-light darkness (Fig. 5b). AtZEP, AtPsbS, SvPsbS lines had reduced NPQ decay time constants, which means faster NPQ decay in the dark (Fig. 7c, d, e). The faster NPQ decay rates in AtZEP lines were consistent with the reduced zeaxanthin levels in these lines after 5 min post-light darkness. Though the faster NPQ decay rates in AtPsbS, SvPsbS lines could not similarly be explained by differences in zeaxanthin content (Fig. 5b), our results are consistent with results reported in Arabidopsis AtPsbS-OE lines which suggested increased PsbS abundance accelerated NPQ relaxation (Steen *et al*., 2020). We also quantified the NPQ decay using the 2^nd^ order exponential decay and got similar results (e.g. AtVPZ-1), though the effects were separated into two phases and the calculated time constants from the 1^st^ phase showed greater relative variations (Fig. S7).

**Figure 7.**
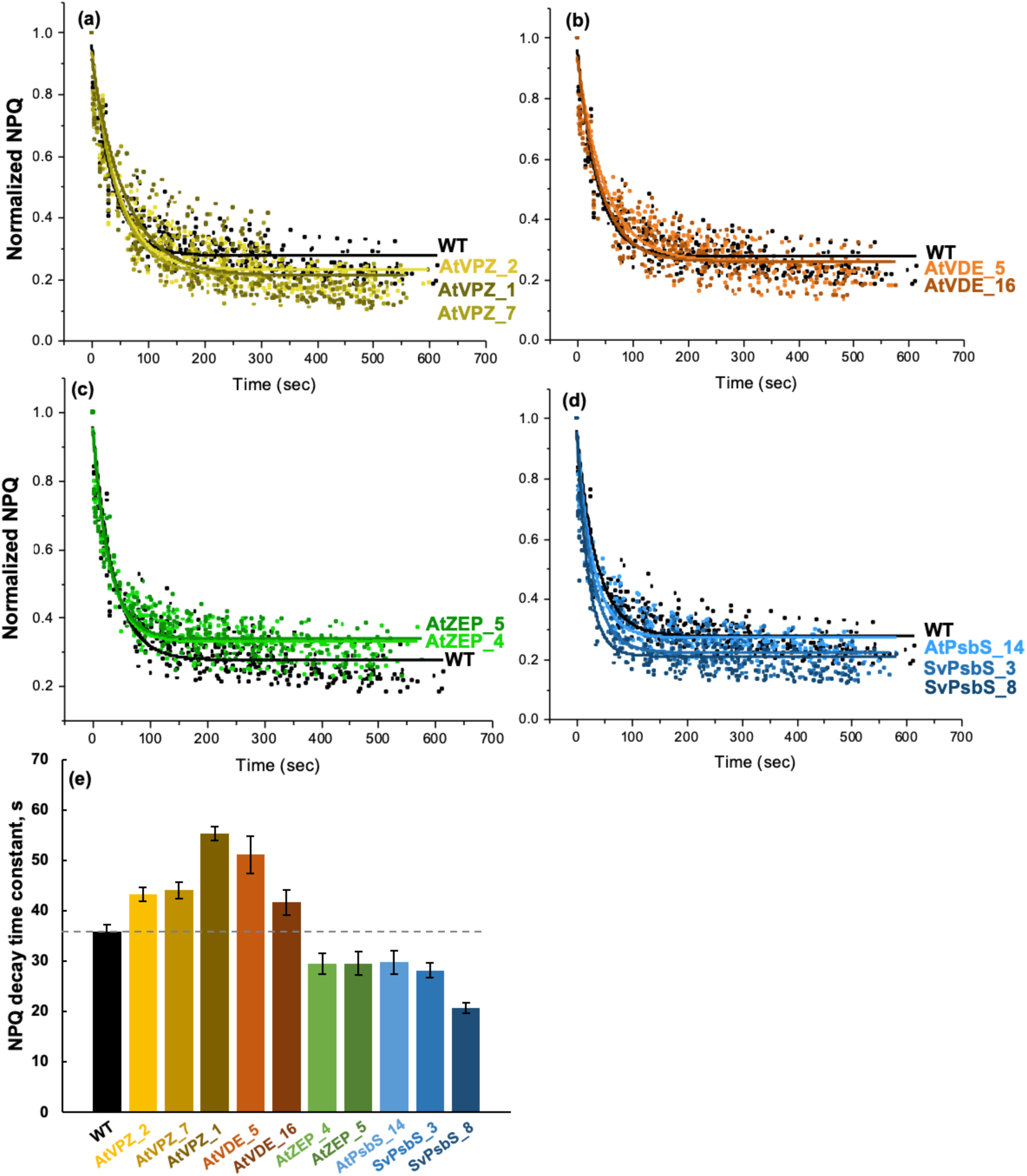
Setaria transgenic AtVPZ, AtVDE lines had slower NPQ decay rates while AtZEP, AtPsbS, SvPsbS lines had faster NPQ decay in dark after the fluctuating light treatment. Plants were treated with fluctuating light as in Figure 6 before NPQ measurement in dark. NPQ data was normalized to the last NPQ values before the post-light dark for each plant. **(a-d)** The NPQ decay curves were fitted using 1^st^ order exponential decay. Genotype names were labeled next to their corresponding curve with the same color as the fitted curve and data points. Note that wild type data has been plotted on each graph for comparison. **(e)** NPQ decay time constant from the 1^st^ order exponential decay fitting for all genotypes. The bigger NPQ decay time constant, the slower NPQ decay rate. The dashed lines mark the WT level. Mean ± SE, n=3-5.

In contrast to fluctuating light, the differences in NPQ capacity between transgenic lines and WT were smaller with gradual increased light (light response curves) (Fig. S8a). But SvPsbS, AtVPZ-1, AtVDE-5 lines still had significantly higher NPQ than WT. AtVDE, AtZEP, and SvPsbS lines often had lower net CO_2_ assimilation rates than WT (significantly lower in SvPsbS-8, AtVDE-5, and AtZEP-5). The AtVPZ-1 line had WT-level net CO_2_ assimilation rates despite higher NPQ (Fig. S8a, c). The differences in PSII operating efficiency from WT were minimal in these transgenic lines (except for SvPsbS-3 and AtVDE-5 with lower values) (Fig. S8b).

In light-adapted leaves, AtVPZ lines had slightly higher proton motive force and higher proton flux rates than WT; while AtVDE, AtZEP, and SvPsbS lines often had reduced proton motive force and PSI active centers as compared to WT, either slightly or significantly (Fig. S9). The results suggest overexpressing single NPQ genes may cause imbalanced or compromised photosynthesis in Setaria.

We further phenotyped these transgenic lines under different environmental conditions (Fig. 8, 9, and S10). These environmental conditions we used affected WT plants at different levels in terms of plant height, wet, or dry biomass (Fig. S10a-c). Overall, the values of wet biomass were highly correlated with those of dry biomass across almost all conditions (Fig. S10d). The AtVPZ lines, especially AtVPZ-1, were taller and/or had more biomass than WT under six out seven conditions we tested: including control, high light, high temperature, greenhouse (August run), drought, and low light conditions (Fig. 8, 9, and S10). AtVPZ-2 and AtVPZ-7 lines had improved growth in three and four conditions based on plant height, wet or dry biomass data (Supplemental Table S1a). The AtVDE lines also had increased dry biomass under control or high temperature conditions (Fig. 8). Additionally, we phenotyped these transgenic lines twice (August 2022 and July 2024) in greenhouse conditions where the environmental parameters were controlled but could be affected by outside ambient conditions (Fig. 9). The August experiment had higher temperatures than the July run, with 5°C differences in maximum high temperatures (Fig. 9a). Our results show that the AtVPZ lines grew better than WT in the August experiment with higher temperatures, but worse than WT in the July experiment with lower temperatures, suggesting some potential links between improved growth and high NPQ/zeaxanthin under high temperature conditions.

**Figure 8.**
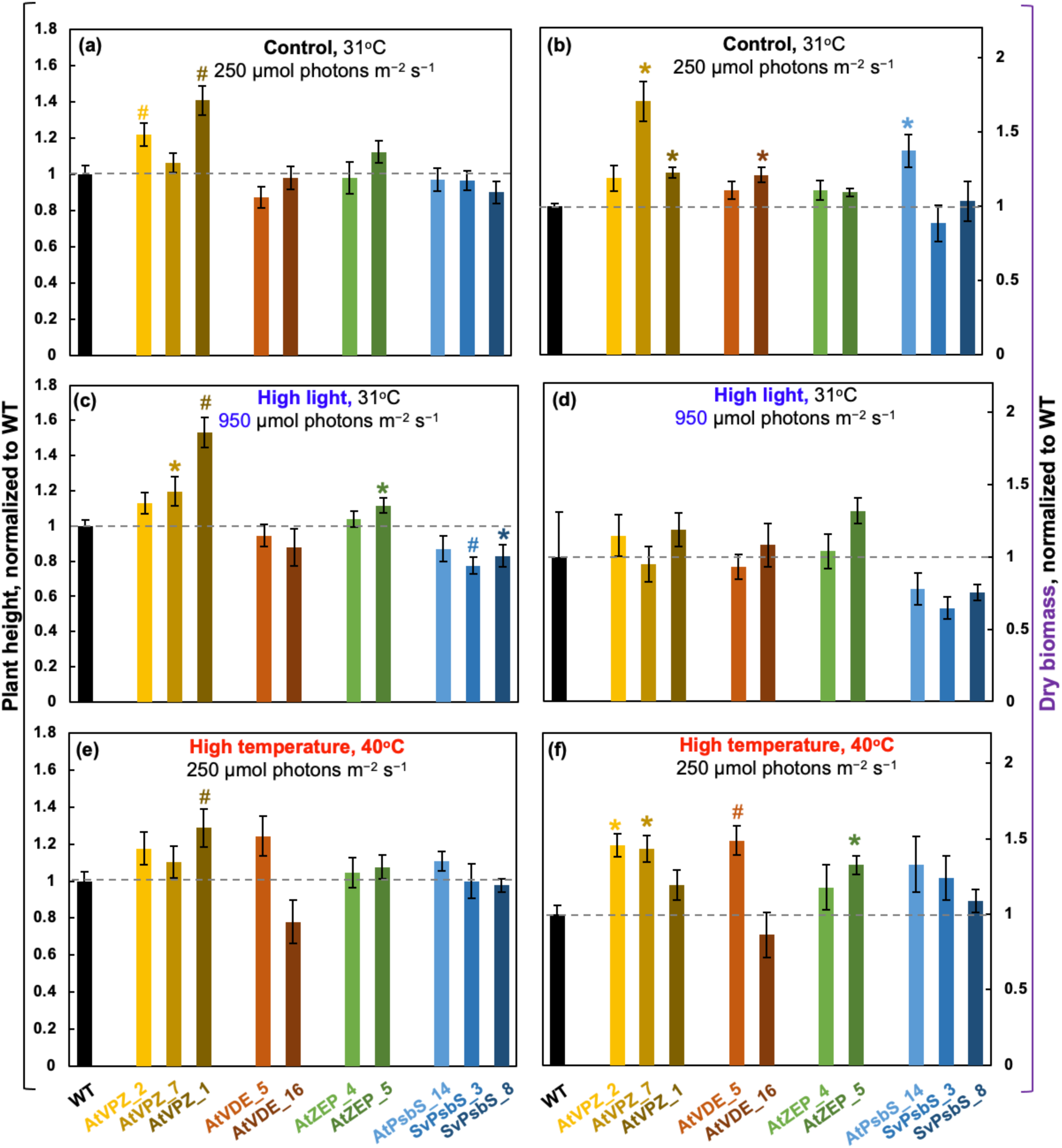
Setaria transgenic AtVPZ lines had increased plant height and/or dry biomass as compared to WT plants under the control, high light, and high temperature conditions. Plant height **(a, c, e)** and whole plant dry biomass **(b, d, f)** under different conditions were quantified and normalized to the mean values of WT plants grown under the same conditions. Plants were grown under the control conditions (constant 31 °C, 250 µmol photons m^−2^ s^−1^ light, 12/12 h day/night) for 9 days before exposed to control **(a, b)**, high light (**c, d,** 950 µmol photons m^−2^ s^−1^ light), or high temperature **(e, f,** constant 40°C) conditions for 5 days. Other environmental parameters of the high light or high temperature conditions stay the same as the control condition. The dashed lines mark WT levels. *, P<0.05; #, P<0.01, transgenic lines were compared to WT under the same condition using a Student’s two-tailed *t*-test assuming unequal variance. Mean ± SE, n=3-33.

**Figure 9.**
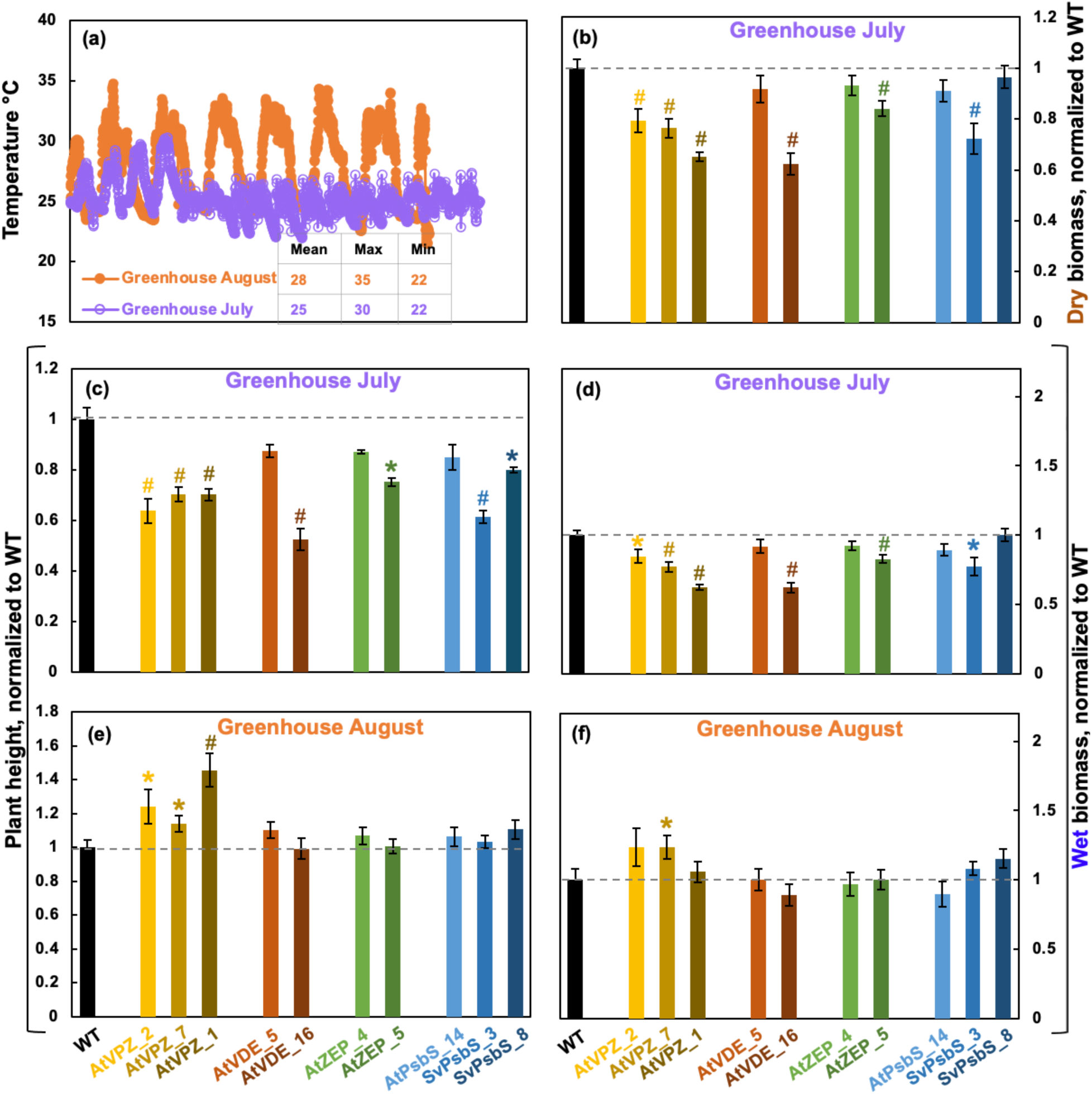
Setaria transgenic AtVPZ lines grew better under warmer temperatures but worse under cooler temperatures than WT in greenhouses. (**a**) Air temperature data of two runs in greenhouses. The August run had warmer temperatures than the July run. The greenhouses had environmental controls, but the inside air temperatures were affected by the outside weather conditions. August run temperature setting: 31/22°C day/night. July run temperature setting: 25/22°C day/night. **(b)** Plant dry biomass in July. The dry biomass data for the August run was unavailable. Plant height **(c, e)** and whole plant wet biomass **(d, f)** after growing in greenhouses. The dashed lines mark WT levels. *, P<0.05; #, P<0.01, transgenic lines were compared to WT under the same condition using a Student’s two-tailed *t*-test assuming unequal variance. Mean ± SE, n=6-14.

The xanthophyll cycle is connected with the ABA pathway as violaxanthin is the precursor of ABA and ZEP is an important enzyme in ABA biosynthesis (Ivanov *et al*., 1995; Ederli *et al*., 1997; Park *et al*., 2008; Jahns *et al*., 2009; Latowski *et al*., 2011; Kaiser *et al*., 2019). We measured the leaf ABA levels in these transgenic lines and WT plants under three selected conditions: control, drought, and high temperatures of 40°C (Fig. S11). In WT, leaf ABA levels increased slightly under drought and high temperatures as compared to the control condition. Overall, there were no differences in ABA levels in these transgenic lines as compared to WT.

## Discussion

NPQ is essential to prevent photodamage in plants under excess light conditions, especially during stressful conditions when photosynthesis is compromised (Rochaix, 2014; Dietz, 2015). NPQ has been well studied in C_3_ plants, but much less so in C_4_ plants, with translation of research complicated by biochemical and anatomical differences between the two photosynthetic pathways (Wang *et al*., 2011; von Caemmerer & Furbank, 2016). Through transient expression in Setaria protoplasts, and characterization of stable transgenic plants, we investigated the function of NPQ genes from the C_3_ model Arabidopsis in the C_4_ model Setaria, demonstrated that overexpression of AtVDE and AtZEP achieved similar results in Setaria as it did in C_3_ plants, revealed the possible incompatibility of AtPsbS with SvPsbS in Setaria, evaluated VPZ strategies in Setaria, and helped expand our functional understanding of NPQ in C_4_ plants.

VDE, ZEP, and PsbS proteins in Setaria are quite similar to those in Arabidopsis, in terms of protein sequence and structure prediction (Fig. S1-3). The phenotypes of transgenic lines overexpressing *AtVDE* and *AtZEP* in Setaria resembled those in C_3_ plants: overexpressing *AtVDE* increased zeaxanthin formation and NPQ induction while overexpressing *AtZEP* decreased zeaxanthin formation and NPQ amplitude (Fig. 1. 5, 6) (Hieber *et al*., 2001; Leonelli *et al*., 2016). These results suggest that the activity of VDE and ZEP, and functional contributions of zeaxanthin to NPQ are similar in C_3_ and C_4_ plants.

The results of *AtPsbS* overexpression in Setaria are surprising, including the significantly reduced NPQ in *AtPsbS*-OE protoplasts (Fig. 1), and the non-Mendelian recovery of homozygous AtPsbS lines (Table 1). The only stable AtPsbS line recovered exhibited one of the highest *AtPsbS* transcript expression levels relative to the reference transcripts but ultimately had WT-level PsbS protein abundance and NPQ (Fig. 3, 4, 6). These results suggest that accumulation of AtPsbS protein in Setaria is costly to photosynthetic efficiency and plant fitness, or it may be largely post-transcriptionally silenced or degraded as we observed in the AtPsbS_14 and AtVPZ lines. These unusual results indicate possible unique regulation of Setaria PsbS.

It is hypothesized that AtPsbS proteins form dimers in dark-adapted leaves where NPQ is not needed; light-induced lumen acidification triggers the monomerization of PsbS dimers, which induce the conformational change of LHCII for NPQ (Bergantino *et al*., 2003; Correa-Galvis *et al*., 2016; Krishnan-Schmieden *et al*., 2021). Our computational analysis showed that AtPsbS protein interacts with SvPsbS protein, with an interaction strength score higher than any other PsbS protein pairs we checked, including any two possible pairs of PsbS from Setaria, Arabidopsis, tobacco, and soybean (Fig. 2c). Thus, the heterologous expression of AtPsbS in WT Setaria protoplasts may affect functional PsbS-dependent protein-protein interactions that negatively affect photosynthesis (Fig. 1). The reduced NPQ capacity of At/Sv-PsbS pair may also contribute to the difficulties in identifying homozygous AtPsbS lines (Table 1).

In tobacco and soybean, overexpression of all three Arabidopsis VPZ genes increased the rate of NPQ induction and relaxation, which was hypothesized to sustain sufficient photoprotection under high light but also support efficient photosynthesis under low light (Kromdijk *et al*., 2016; De Souza *et al*., 2022). We employed three Setaria AtVPZ lines which overexpressed all three NPQ-related genes from Arabidopsis in Setaria (Stone *et al*., 2024). The AtVPZ lines had improved growth in 3-6 conditions (Fig. 8-9, S10, Supplemental Table S1a), however they had increased zeaxanthin, faster NPQ induction, but slower NPQ relaxation, which correlated with their higher zeaxanthin levels than WT in 5 min post-light dark (Fig. 5-7). This is especially true for the AtVPZ-1 line, which had the highest NPQ induction but slowest NPQ relaxation among all three AtVPZ lines and grew better in six out of the seven conditions we tested. Our results suggest that overexpressing all three *AtNPQ* genes can improve C_4_ plant growth, but the faster NPQ relaxation may be dispensable or less critical for growth improvement in C_4_ plants under stressful conditions.

It has been proposed that the effects of VPZ strategy on plant growth improvement may be species-specific and the stoichiometries rather than the absolute abundances of the three NPQ proteins are important (Croce *et al*., 2024). For example, the YZ-26-1C soybean line had little AtPsbS protein and the lowest abundance of AtZEP and AtVDE proteins as compared to other transgenic lines, but it still had 21.7% yield increase as compared to WT plants, while two other soybean lines (ND-17-20 and ND-19-8A) had significantly increased all three AtNPQ proteins but had no growth improvement (De Souza *et al*., 2022). We also performed stoichiometry analysis of NPQ proteins in our work and published literature (Kromdijk *et al*., 2016; Garcia-Molina & Leister, 2020; De Souza *et al*., 2022; Lehretz *et al*., 2022) (Supplemental Table S1a). But due to the limited sample size in different species and difference in protein quantification, there seemed to be no clear patterns associating ideal stoichiometry of NPQ proteins and plant growth improvement. However, our results suggest that growth improvement could be achieved without substantially higher amounts of PsbS proteins. Given the fitness gains despite WT-like levels of AtPsbS proteins in our AtVPZ lines, the simplified VZ strategy (instead of VPZ) may make it easier for genetic engineering to improve plant growth.

Additionally, our photosynthesis measurements showed that overexpressing single NPQ genes may cause imbalanced or compromised photosynthesis in Setaria, in terms of photosynthetic light reactions and carbon fixation (Fig. S8, S9). Altogether, our data support the idea that awareness of native photoprotective capacities and needs, alongside investigation in physiologically relevant environments, will be critical in engineering increased photosynthetic efficiency at the leaf and field scales (Croce et al., 2024).

Furthermore, our research reveals potential links among increased zeaxanthin, NPQ, and thermotolerance in C_4_ plants. Our greenhouse experiments show that AtVPZ lines grew better under warmer temperatures but worse under cooler temperatures than WT (Fig. 9). The increased zeaxanthin and NPQ in our AtVPZ lines may help stabilize thylakoid membranes and protect photosynthetic damages under warm temperatures (Sharkey & Zhang, 2010; Demmig-Adams *et al*., 2020; Anderson *et al*., 2021). This is further supported by the improved growth of AtVPZ lines and AtVDE-5 lines under high temperature of 40°C (Fig. 8). In agreement with this, a recent result showed that overexpression of VDE alone increased NPQ induction and biomass production by about 11-16% in rice under the field conditions (Xin *et al*., 2023). These rice plants were grown during the summertime in southern China (Songjiang), with daily average temperature likely above 27°C. Our Setaria plants were grown in growth chambers with a control temperature of 31°C.

Our research advances the understanding of the regulation of NPQ in C_4_ plants and paves the way to improve C_4_ photosynthesis. Future exciting questions regarding NPQ regulation in C_4_ plants may include: (1) Does NPQ have cell-type specificity (M or BS cells) in C_4_ plants? (2) How is PsbS regulated to achieve sufficient photoprotection and efficient photosynthesis in C_4_ plants? (3) How do M and BS cells coordinate to regulate NPQ and photosynthesis? (4) How could we further improve C_4_ photosynthesis under stressful conditions? We hope our results will cultivate more interest in C_4_ NPQ and promote research to answer these important and intriguing questions.

## Acknowledgements

The research was supported by the Defense Advanced Research Projects Agency (DARPA) (HR001118C0137 to RZ and DAN) and start-up funding from Donald Danforth Plant Science Center (DDPSC, to RZ). This work to generate the Setaria AtVPZ lines was supported by the DARPA Advanced Plant Technologies program to TJL and MAG under contract HR001118C0146. DPT was supported by the Berkeley Fellowship and the NSF Graduate Research Fellowship Program (Grant DGE 1752814). KKN is an investigator of the HHMI. The authors acknowledge the use of the metabolite profiling facility of the Bindley Bioscience Center, a core facility of the NIH-funded Indiana Clinical and Translational Sciences Institute, and an National Science Foundation (NSF) grant (IOS-2140119 to SM), Oak Spring Garden Foundation’s Fellowship in Plant Science Research to CNK for aiding in the quantification of ABA levels. Some bioinformatics data analysis was supported by an NIH grant (R01GM093123) and two NSF grants (DBI2308699 and CCF2343612) to JC. GJ received funding through NSF award 2019516. We thank Drs. Stephen Long, Johannes Kromdijk, Katarzyna Glowacka, Alizée Malnoë, and Ivan Baxter for discussing some of our results, and Drs. Dong-Yeon Lee and Margaret Wilson for suggestions on our Golden Gate Cloning and protoplast isolation. We appreciate the help from DDPSC Plant Transformation Facility for generating the AtNPQ and SvPsbS transgenic lines and DDPSC Integrated Plant Growth Facility for assistance with plant growth. We thank Dr. Joyce Van Eck and the Boyce Thompson Institute Center for Plant Biotechnology Research for generating the Setaria AtVPZ transgenic plants and their assistance with growth protocols.

## Disclaimer

The views, opinions and/or findings expressed are those of the author and should not be interpreted as representing the official views or policies of the Department of Defense or the U.S. Government. Distribution Statement “A” (Approved for Public Release, Distribution Unlimited).

## Author contributions

RZ designed and supervised the whole project and wrote the initial paper draft. GM screened and identified most of the Setaria AtNPQ transgenic lines, optimized most LICOR and phenotyping measurements, analyzed all the photosynthetic and phenotyping data, and led RT-qPCR analysis, prepared most of the figures, and drafted the method section. CMM optimized the protoplast isolation and transformation protocol, generated constructs to make AtNPQ and SvPsbS lines, and screened T_o_ of these lines. EK performed some of the LICOR and multispeQ experiments, sample harvesting, chlorophyll extraction, stress treatments, and helped with some of the RT-qPCR analysis. SP, KH, EK performed western blots. DPT, KKN performed HPLC analysis of pigments. WEM optimized chlorophyll fluorescence measurements in Setaria protoplasts and EB performed the measurements. CB helped with some of the plant harvesting and LICOR measurements. DARPA LISTENS team genotyped and identified the AtVPZ lines: LAG (genotyped these lines under supervision of MAG), GJ (interpreted early generation expression and genotyping results), PMT (identified transgene insertions), WDS (performed initial characterization of these lines under supervision of TJL) and XJK (performed cloning and design under supervision of FZ). FZ also provided the backbone of AtNPQ constructs. DAN helped optimize the protoplast isolation/transformation protocol and coordinated the collaboration among DARPA teams. JL and JLC performed computational modeling of NPQ proteins. DHP performed part of the greenhouse and some of the LICOR experiments and statistical analysis, and helped polish some of the figures. JS helped with some of the statistical analysis. SAMMK and CNK quantified leaf ABA levels. RZ, DPT, KKN, DHP, GM, MAG, JL, GJ, EK, SAMM, and KH helped revise the manuscript.

## Data availability statement

All data that supports the findings of this study are available within the paper and within its supplemental materials published online.

---

**Supplemental Table S1.** (**a**) NPQ protein stoichiometries; (**b**) Primers used.

**Supplemental File S1**. Protein structure and interaction prediction

## Supplemental figures

**Figure S1.**
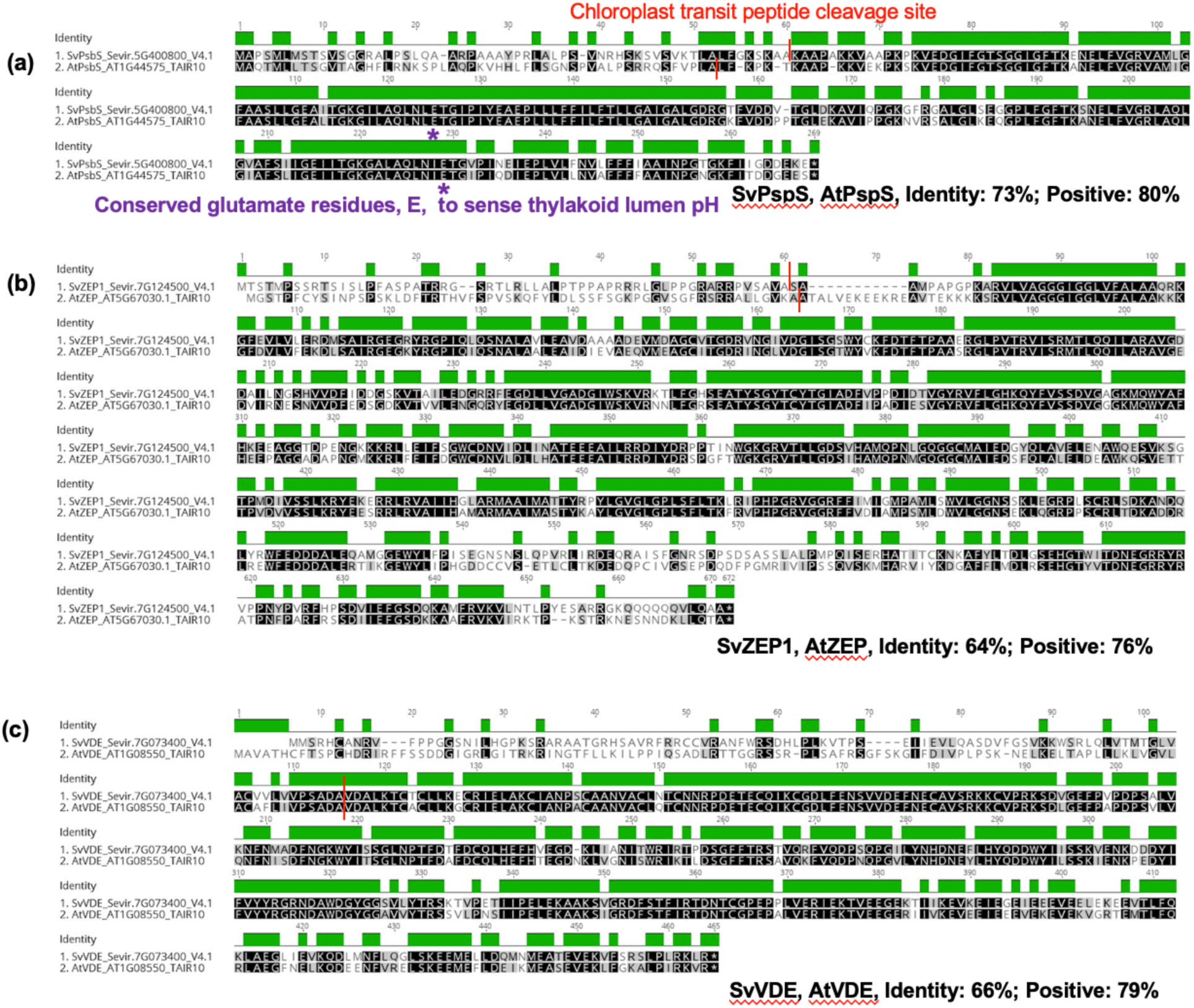
Protein alignment of NPQ proteins from Arabidopsis and Setaria. Alignment of Setaria and Arabidopsis PsbS **(a)**, ZEP **(b)**, and VDE **(c)** proteins. Arabidopsis and Setaria each have one copy of PsbS and VDE. Arabidopsis has one copy of ZEP while Setaria has two copies of ZEP. The more abundant SvZEP1 was used. Setaria protein sequences were from *Setaria viridis* genome V4.1 in Phytozome. Arabidopsis protein sequences were from TAIR 10 in Phytozome. The purple stars (*) in panel (**a**) denote the conserved glutamate residues of PsbS that sense pH. Chloroplast transit peptide cleavage sites are marked with red vertical lines.

**Figure S2.**
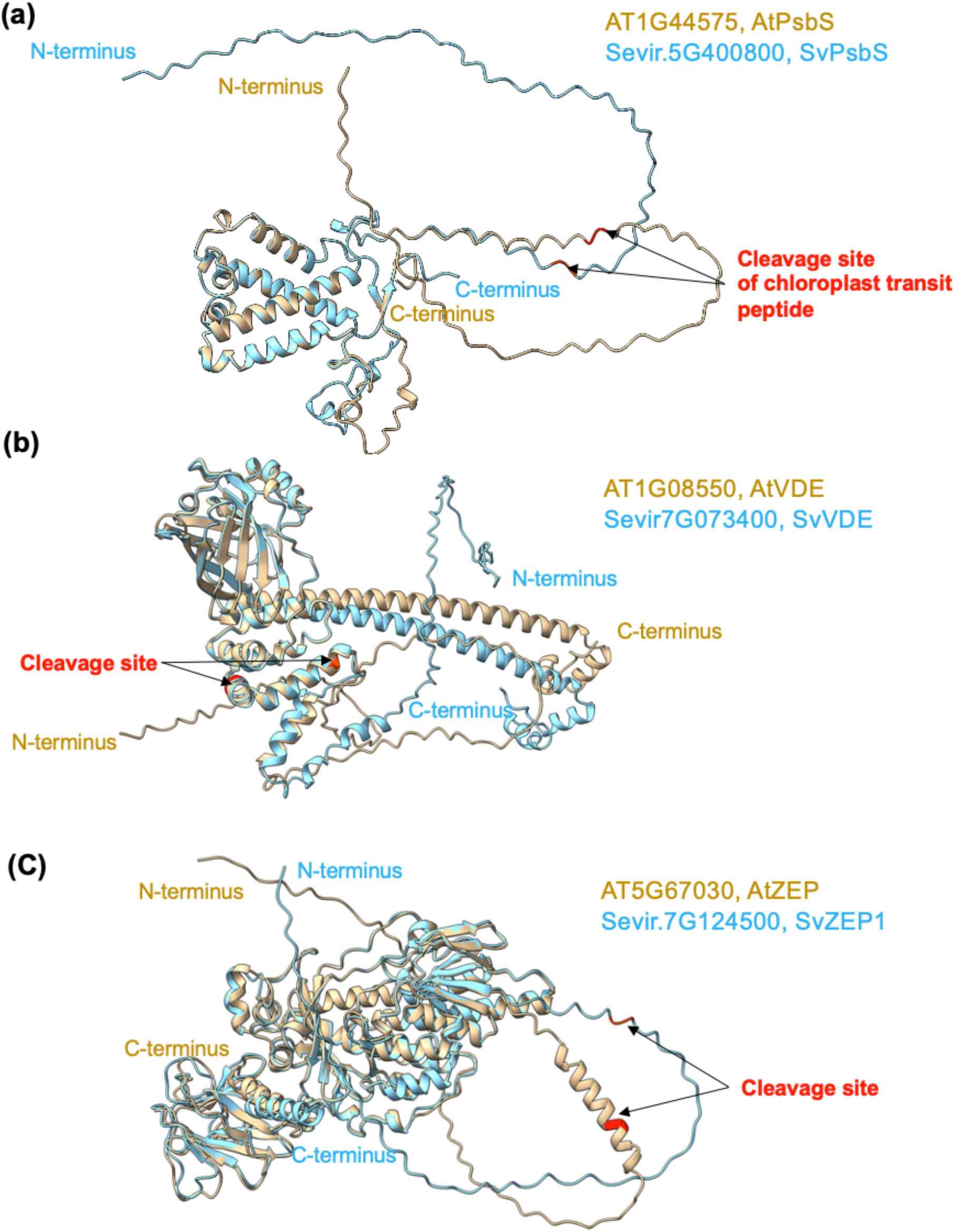
The tertiary structure of Arabidopsis and Setaria NPQ proteins were predicted by MULTICOM3. At, proteins from Arabidopsis. Sv, proteins from Setaria. **(a)** AtPsbS and Sv PsbS; **(b)** AtVDE and SvVDE; **(c)** AtZEP and SvZEP1. The predicted cleavage site for chloroplast transit peptide of each protein is highlighted in red. The N and C terminus of each protein are marked.

**Figure S3.**
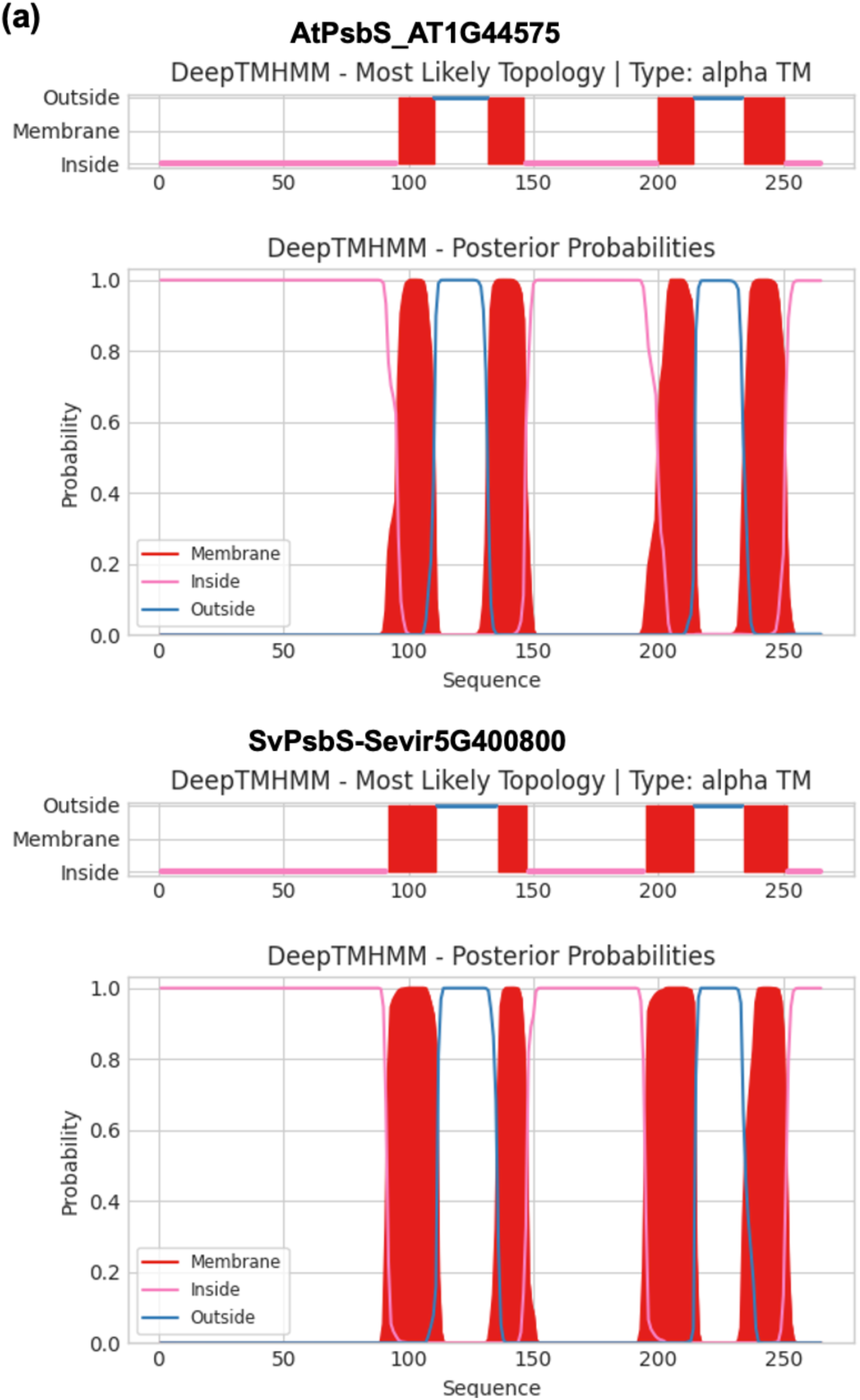
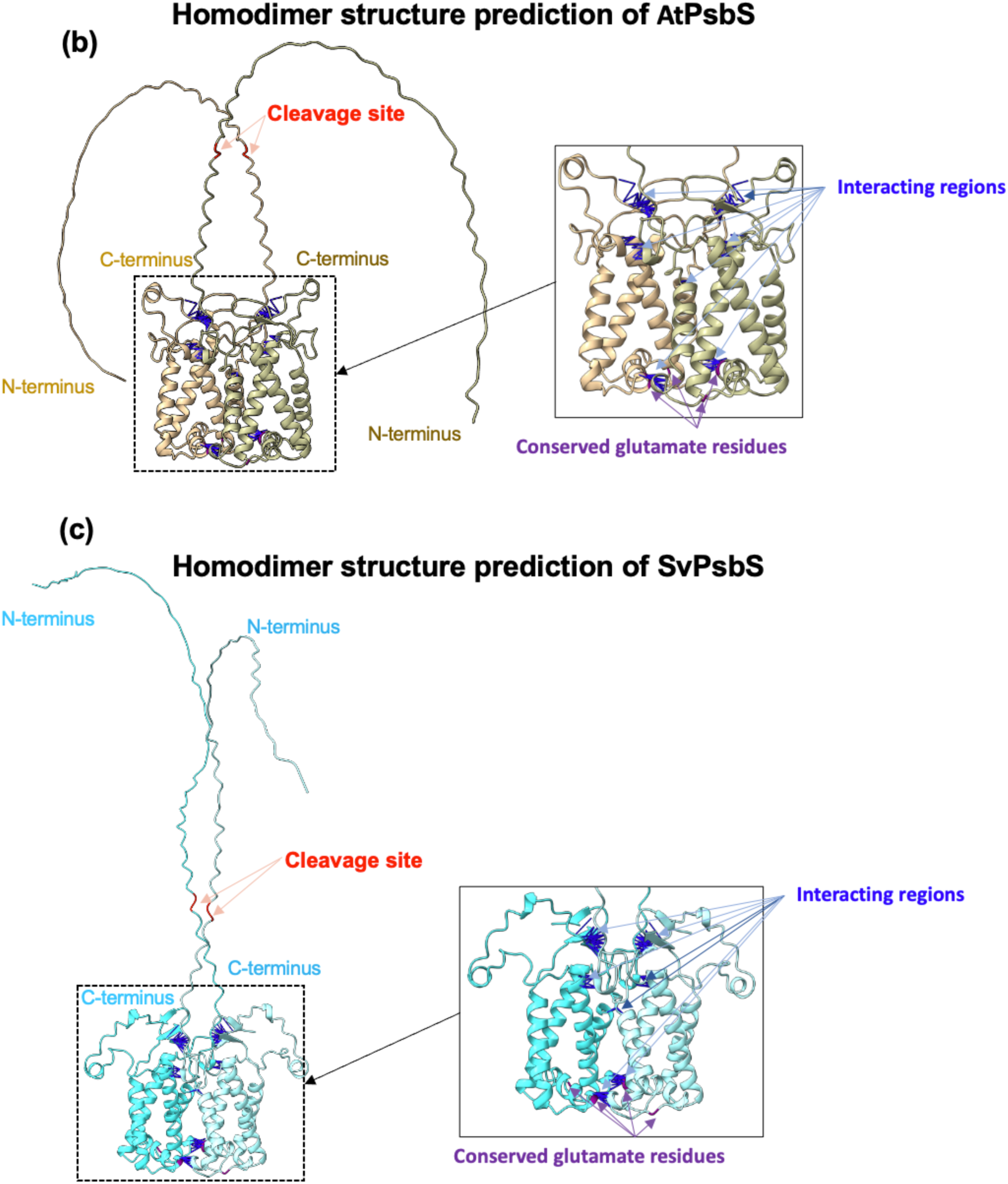
The structures and interactions of Arabidopsis and Setaria PsbS proteins were predicted. **(a)** Arabidopsis and Setaria PsbS proteins both have four transmembrane domains predicted by DeepTMHMM. **(b, c) The dimer structure predictions of At/AtPsbS and Sv/SvPsbS pairs.** AtPsbS (brown) and SvPsbS (cyan) are predicted to self-interact to form homodimers. The predicted cleavage site for chloroplast transit peptide of each protein is highlighted in red. The N and C terminus of each protein are marked. The predicted interacting regions are marked by blue lines. Due to the visualization angle of the predicted structure, only some interaction clusters are shown. A close view of interaction sites is also shown. The conserved glutamate residues of PsbS that sense pH are marked in purple.

**Figure S4.**
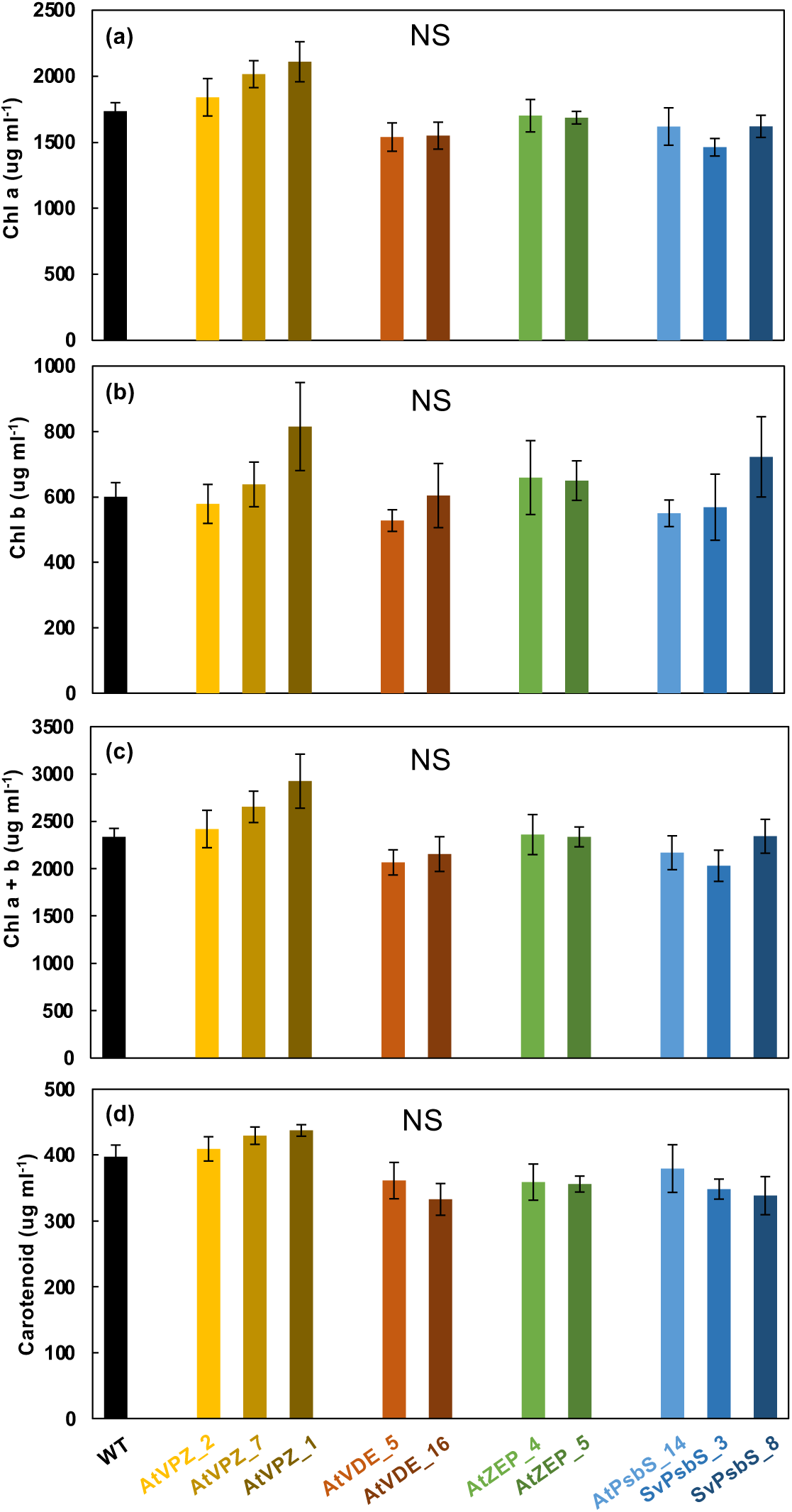
Setaria transgenic lines had no significant differences in chlorophyll or carotenoid contents as compared to WT under the control condition. **(a)** Chlorophyll (Chl) a content. **(b)** Chl b content. **(c)** Chl a and b, or total Chl content. **(d)** Carotenoid content. Transgenic lines were compared to WT using a Student’s two-tailed *t*-test assuming unequal variance. NS, not significant. Mean ± SE, n=3.

**Figure S5.**
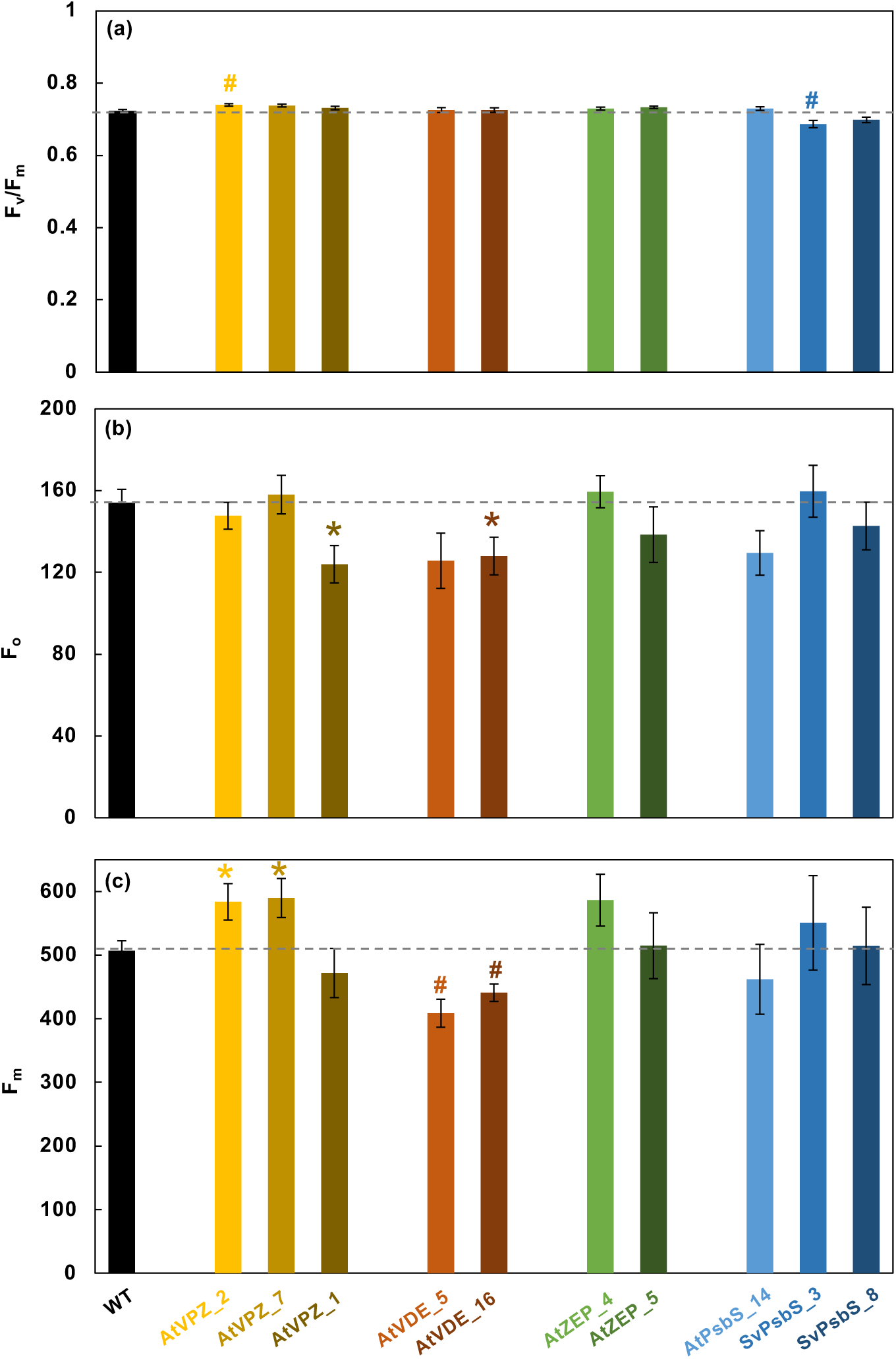
Photosynthetic parameters from dark-adapted Setaria WT and transgenic lines. Plants were dark-adapted for 25 min before measurements. A fully expanded intact 4^th^ leaf was used for chlorophyll fluorescence measurements in a LI-6800 machine. **(a)** Maximum PSII efficiency (F_v_/F_m_), **(b)** minimum and **(c)** maximum chlorophyll fluorescence (F_m_ and F_o_) in dark-adapted leaves. The dashed lines mark WT levels. *, P<0.05; #, P<0.01, transgenic lines were compared to WT under the same condition using a Student’s two-tailed *t*-test assuming unequal variance. Mean ± SE, n=5.

**Figure S6.**
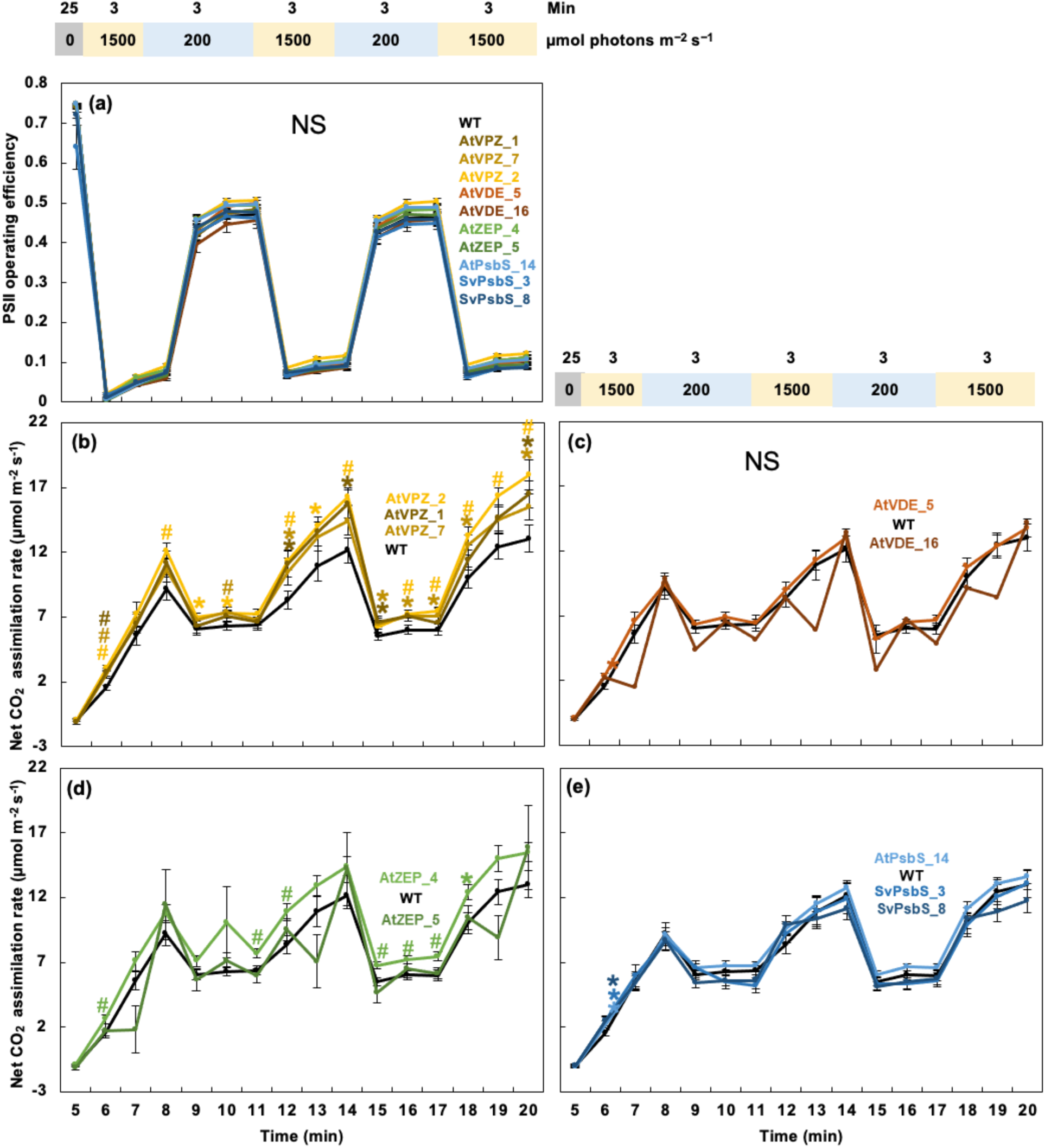
Photosynthetic parameters from Setaria WT and transgenic lines with fluctuating light treatment. Leaves were dark-adapted for 25 min before fluctuating light treatments. **(a)** PSII operating efficiency, **(b-e)** net CO_2_ assimilation rate of all lines during the fluctuating light condition. Genotype names were labeled by the corresponding curves with the same color and order as the curves. *, P<0.05; #, P<0.01, transgenic lines were compared to WT under the same condition using a Student’s two-tailed *t*-test assuming unequal variance. NS, not significant. Mean ± SE, n=18-23.

**Figure S7.**
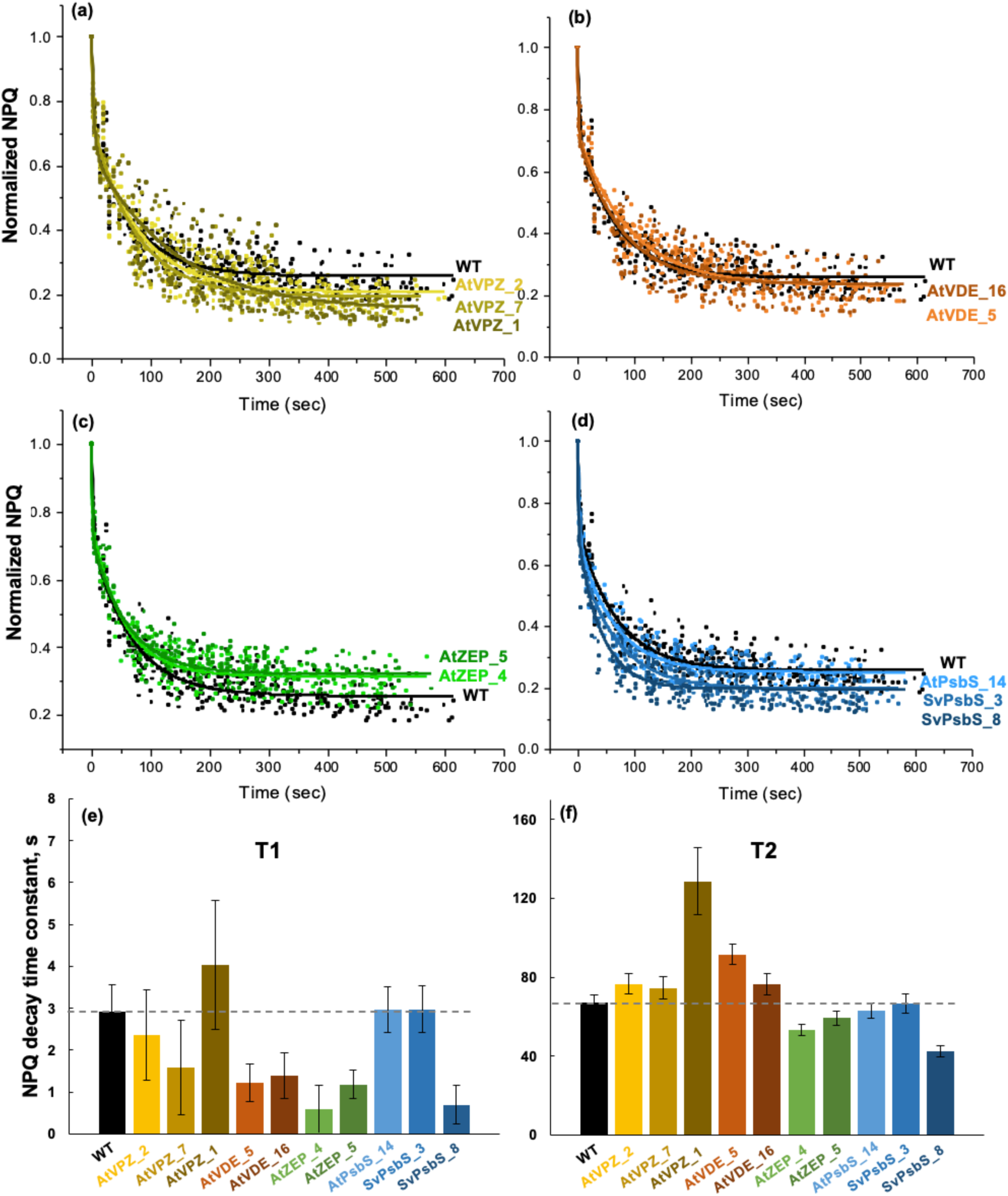
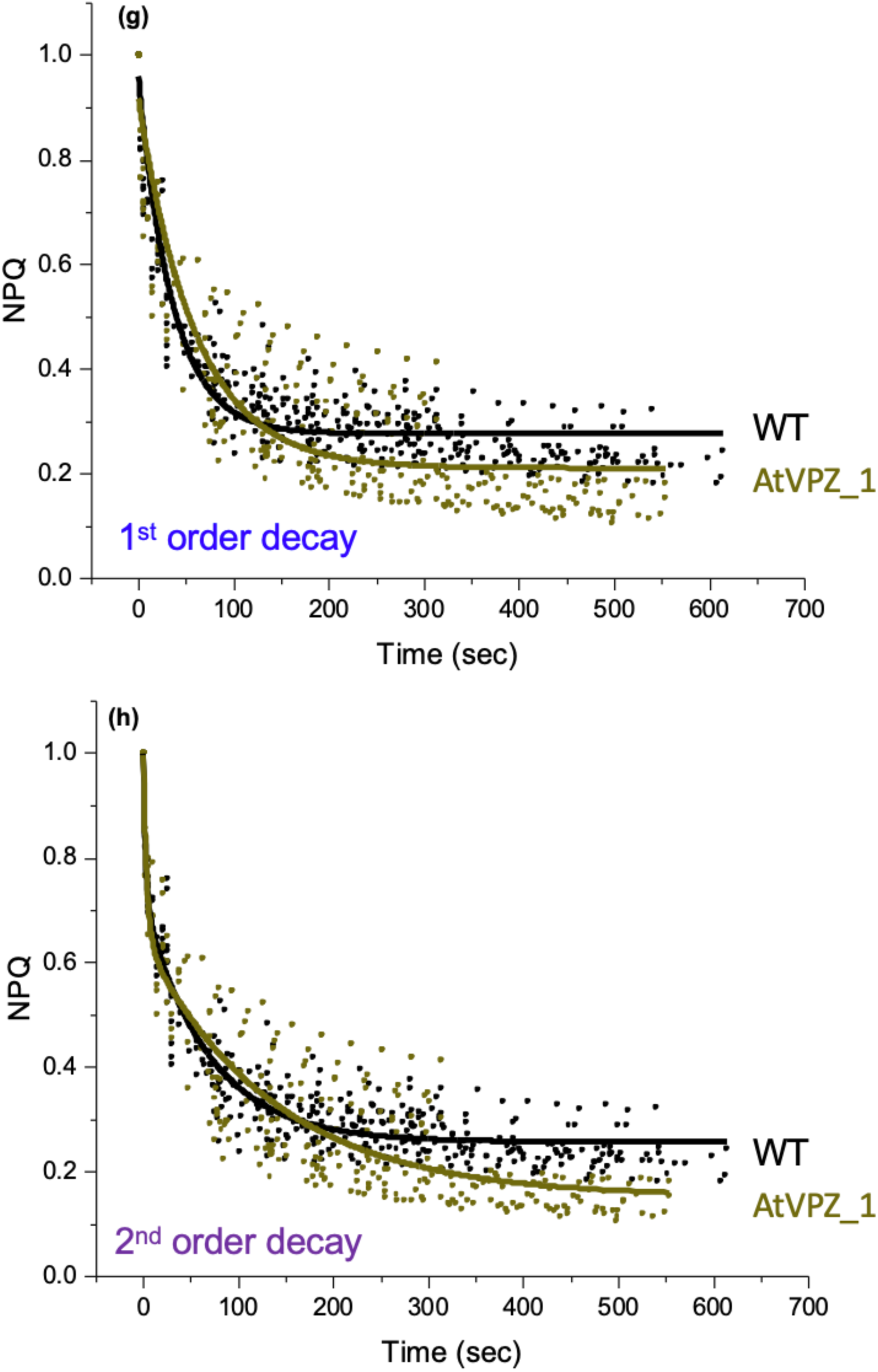
NPQ decay time constants in Setaria WT and transgenic lines were quantified using 2^nd^ order exponential decay. Plants were treated with fluctuating light as in Figure S6 before measurement of NPQ decay in dark. NPQ data was normalized to the last NPQ values before the post-light dark for each plant. **(a-d)** The NPQ decay curves were fitted using 2^nd^ order exponential decay. Genotype names were labeled next to the corresponding curve with the same color as the fitted curve and data points. **(e, f)** NPQ decay time constants from the 2^nd^ order exponential decay fitting for all genotypes, T1 is the time constant for the 1^st^ and faster decay component, T2 is the time constant for the 2^nd^ and slower decay component. Mean ± SE, n=3-5. **(g, h)** The transgenic AtVPZ_1 had slower NPQ decay than WT, evaluated using both 1^st^ and 2^nd^ order exponential decays.

**Figure S8.**
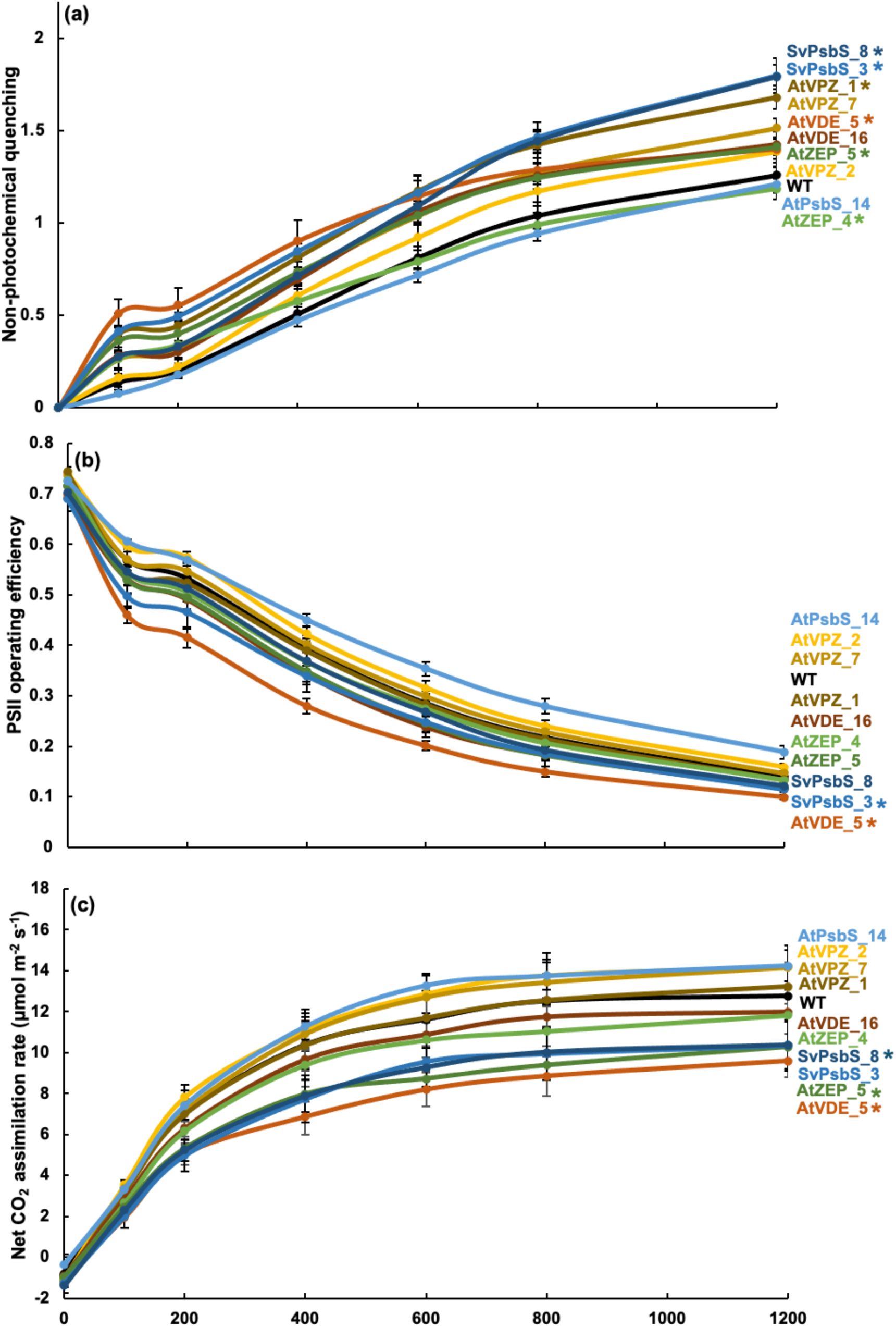
Photosynthetic parameters from Setaria WT and transgenic lines with gradually increased light intensity. **(a)** Non-photochemical quenching, NPQ, **(b)** PSII operating efficiency, **(c)** net CO_2_ assimilation rates were measured in plants with 25 min dark adaptation followed by gradually increased light from 0 to 1500 µmol photons m^−2^ s^−1^. Each light lasted 5 min. Genotype names were labeled by the corresponding curves with the same color and order as the curves. The light response curves of transgenic lines were compared to that of WT using statistical modeling and posterior probabilities, *, P>0.95 as significance. Mean ± SE, n=5-10.

**Figure S9.**
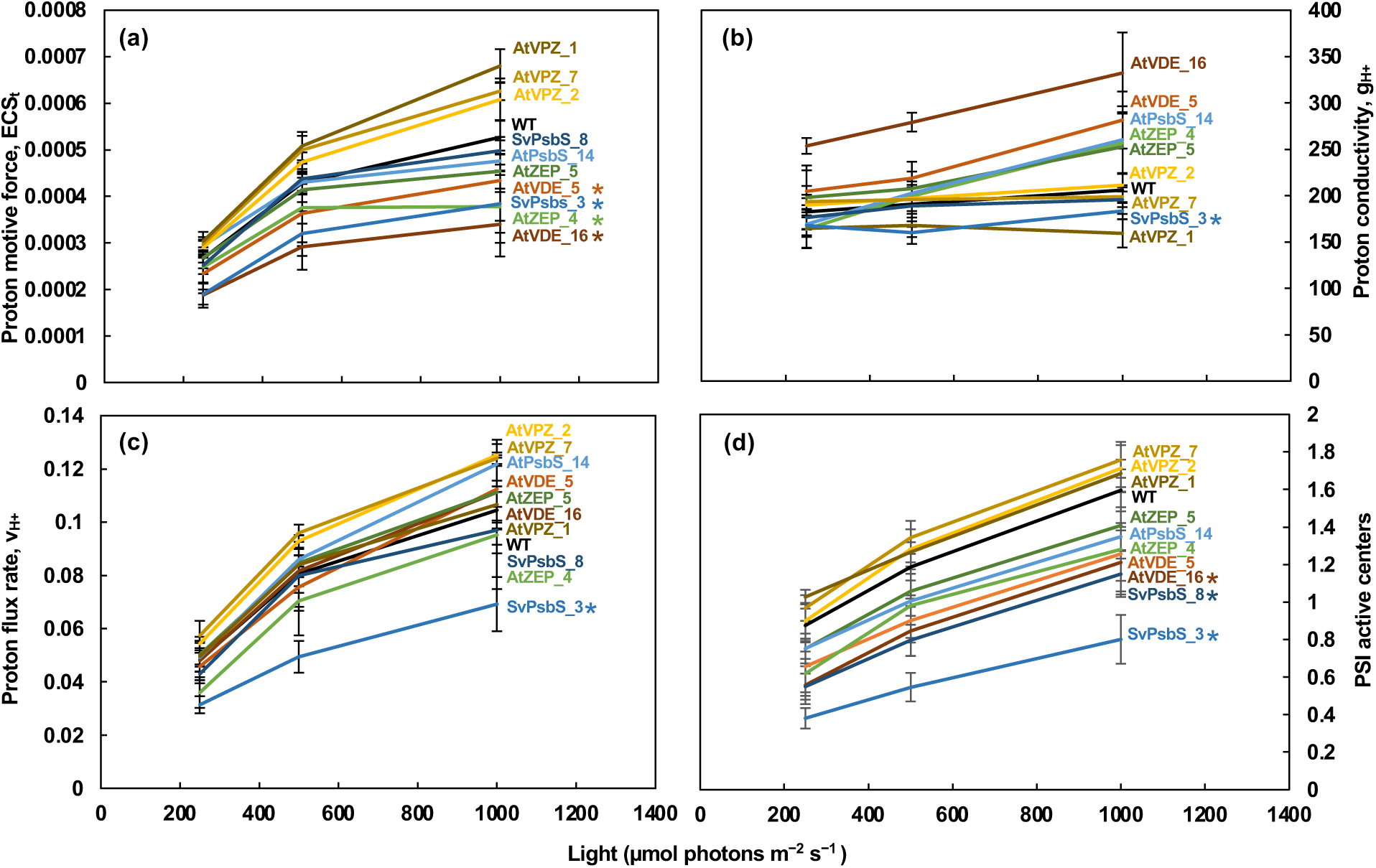
Setaria transgenic SvPsbS lines had reduced fraction of PSI active centers. Photosynthetic parameters were measured in intact leaves of light adapted Setaria WT and transgenic plants using the MultispeQ. **(a)** Proton motive force, ECS_t_, measured by electrochromic shift (ECS), proportional to the transthylakoid proton motive force. **(b)** Proton conductivity, (ɡ_H_^+^ = 1/−_ECS_), proton permeability of the thylakoid membrane and largely dependent on the activity of ATP synthase, inversely proportional to the decay time constant (−_ECS_) of light-dark-transition-induced ECS signal. **(c)** Proton flux rate, v_H+,_ calculated by ECS_t_/−_ECS_, the initial decay rate of the ECS signal during the light-dark transition and proportional to proton efflux through ATP synthase to make ATP. **(d)** The fraction of PSI active centers. The light response curves of transgenic lines were compared to that of WT using statistical modeling and posterior probabilities, *, P>0.95 as significance. Mean ± SE, n=5-10.

**Figure S10.**
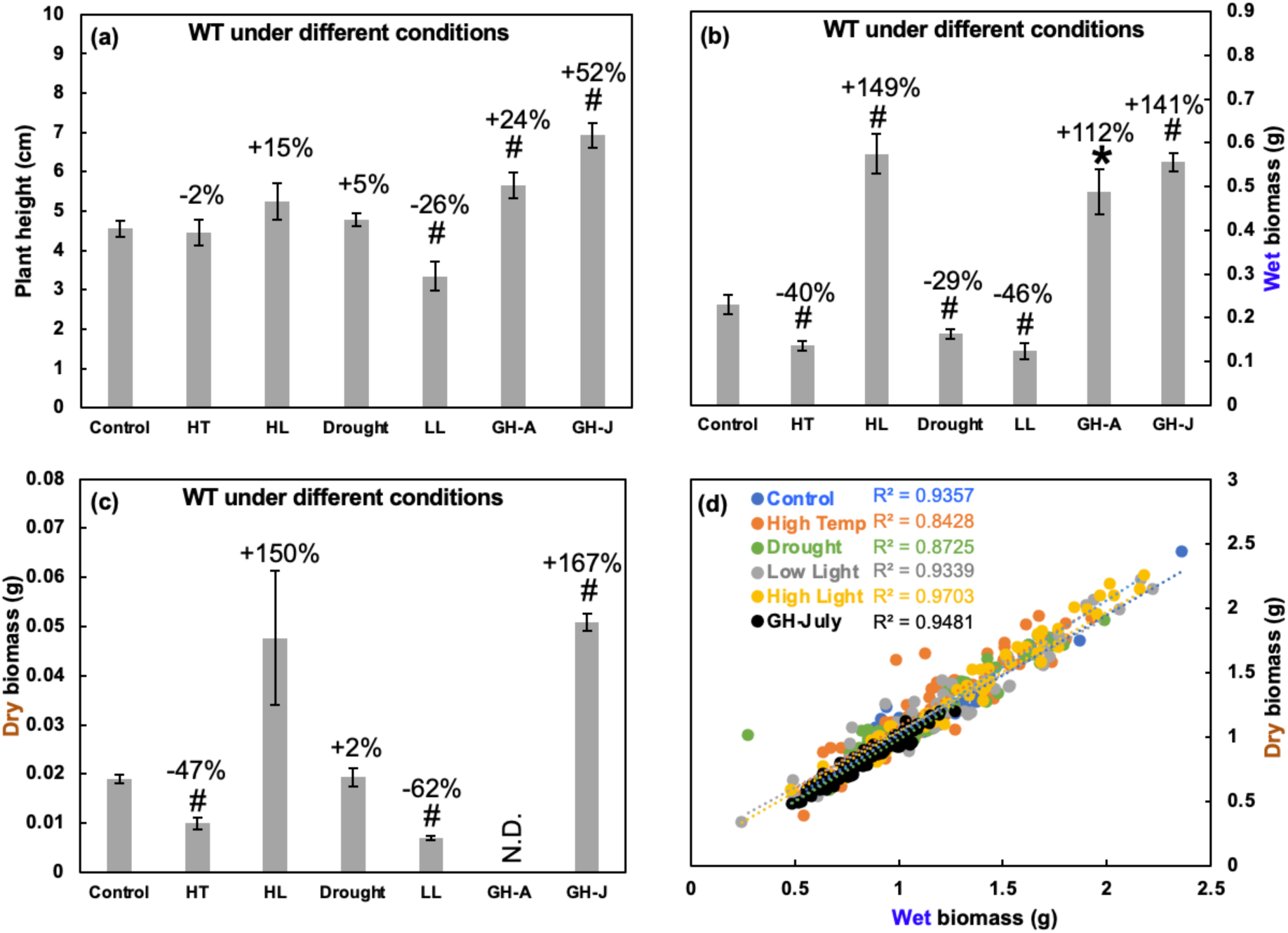
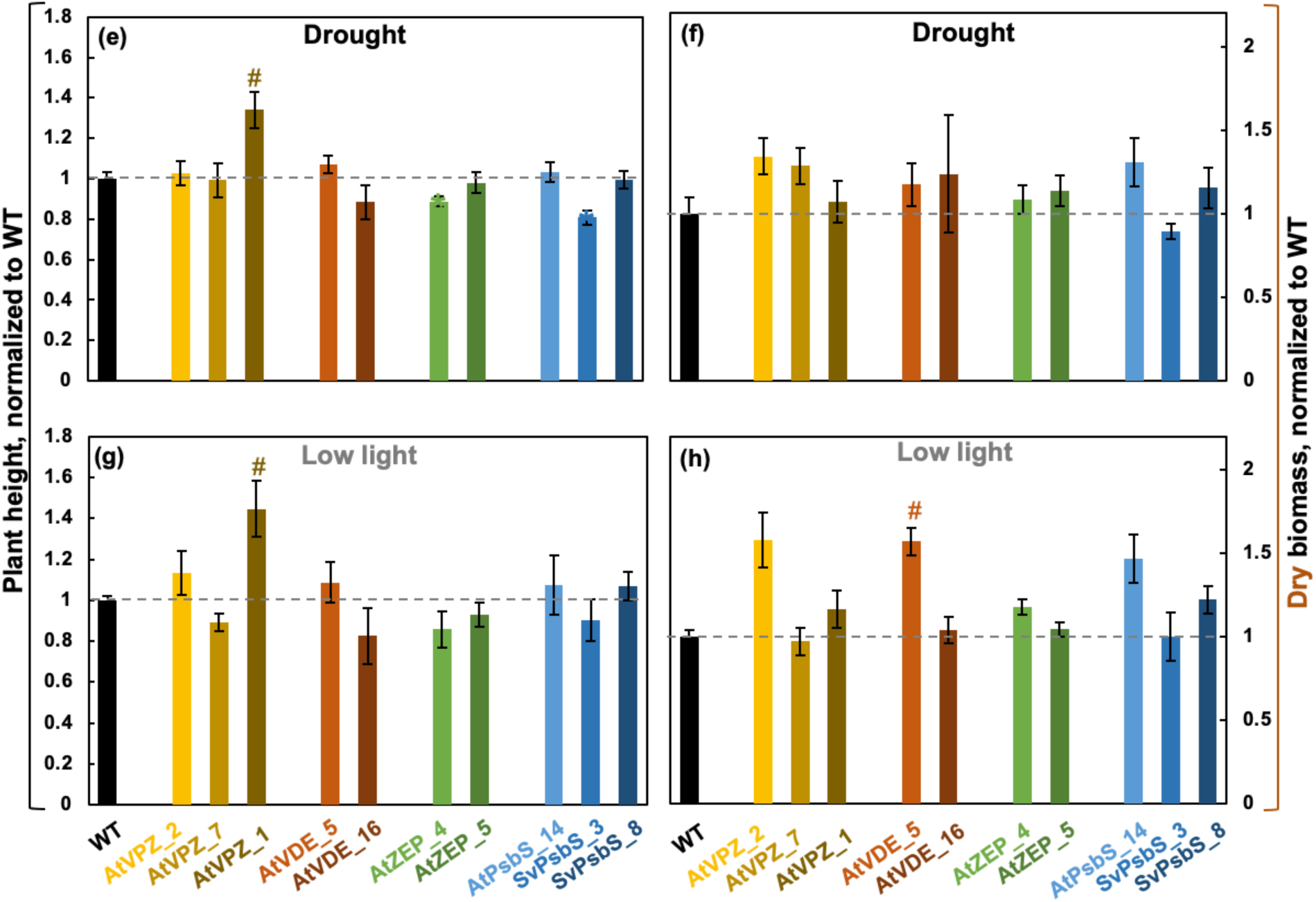
Setaria WT and transgenic lines were phenotyped under various conditions. **(a)** Plant height, **(b)** whole plant wet biomass**, (c)** whole plant dry biomass of WT plants under the control, high temperature (HT), high light (HL), drought, low light (LL), and greenhouse (GH) conditions. Two rounds of greenhouse experiments were conducted, one in August 2022 (GH-A) and the other in July 2024 (GH-J). Plants were grown under the control conditions (constant 31 °C, 250 µmol photons m^−2^ s^−1^ light, 12/12 h day/night, with daily water) for 9 days before exposure to different conditions for 5 days: HT, constant 40°C; HL, 950 µmol photons m^−2^ s^−1^ light; drought, no water; LL, 100 µmol photons m^−2^ s^−1^ light; GH, affected by dynamic changes of natural light and temperatures; other unmentioned environmental parameters stay the same as the control condition. **(d)** Whole plant wet and dry biomass were highly correlated with each other under different conditions. Plant height **(e, g)** and whole plant dry biomass **(f, h)** under different conditions were quantified and normalized to the mean values of WT plants grown under the same conditions. (**e-h**) The dashed lines mark WT levels. *, P<0.05; #, P<0.01, transgenic lines were compared to WT under the same condition using a Student’s two-tailed *t*-test assuming unequal variance. Mean ± SE, n=5-20.

**Figure S11.**
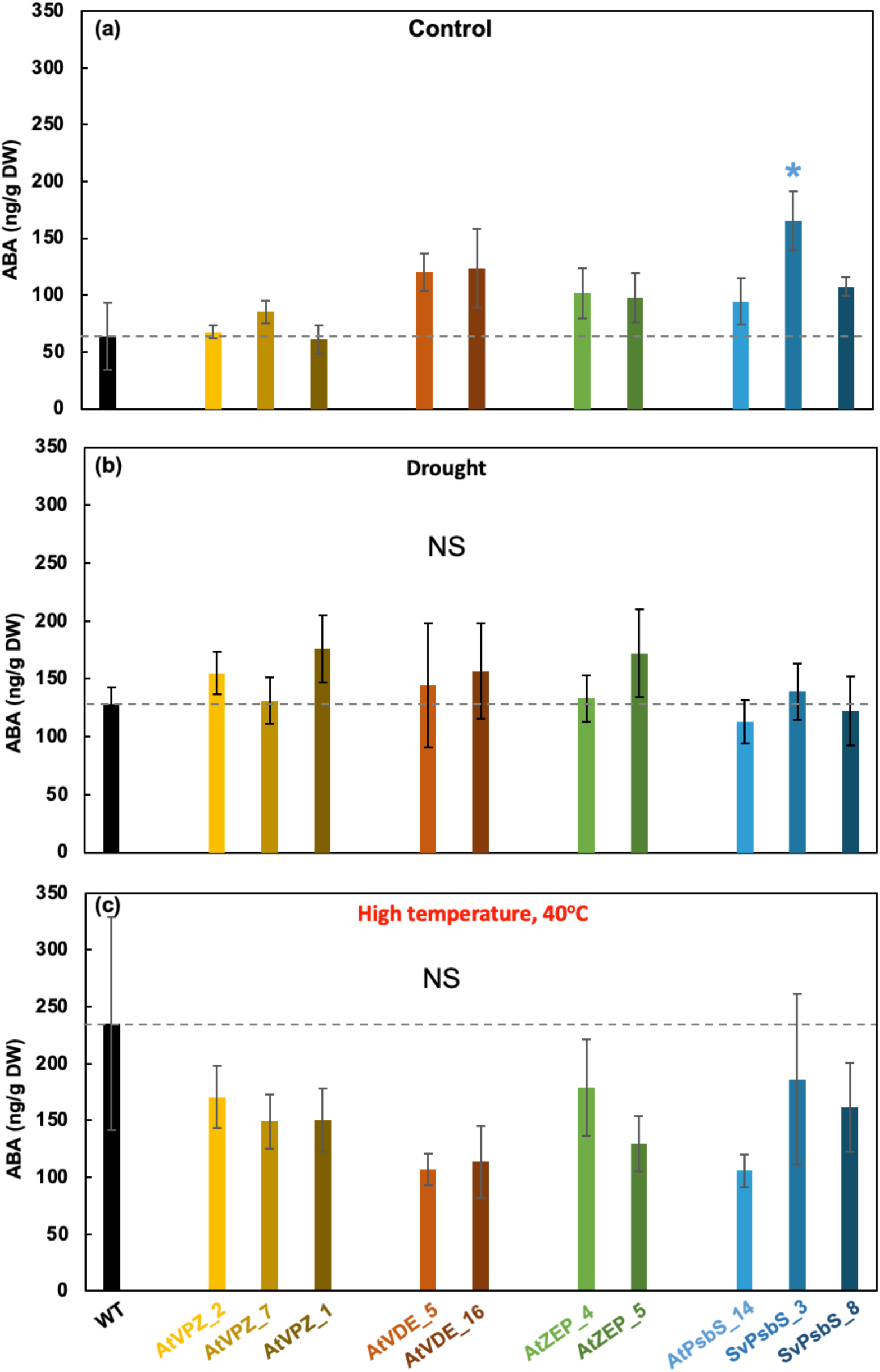
The leaf ABA contents were not significantly different between WT and transgenic lines under the control, drought, and high temperature conditions. Plants were treated with the indicated condition as in Figure 8 and S10 and a fully expanded 4^th^ leaf from a plant after 5-day control or stress treatments was used for ABA measurement. The dashed lines mark WT levels. DW, dry weight of a leaf. Transgenic lines were compared to WT under the same condition using a Student’s two-tailed *t*-test assuming unequal variance. NS, not significant, P>0.05. Mean ± SE, n=4-7.

