## Supplemental_File 1 for "Modification of Non-photochemical Quenching Pathways in the C_4_ Model Plant *Setaria viridis* Revealed Shared and Unique Photoprotection Mechanisms as Compared to C_3_ Plants"

**Supplemental File 1: Protein structure and interaction prediction**

**Part 1:**

**For supplemental Figure S2: Structure prediction of Arabidopsis and Setaria NPQ proteins**

AT1G44575, AtPsbS

Sequence:

MAQTMLLTSGVTAGHFLRNKSPLAQPKVHHLFLSGNSPVALPSRRQSFVPLALFKPKTKAAPKKVEKPKSKVEDGIFGTSGGIGFTKANELFVGRVAMIGFAASLLGEALTGKGILAQLNL**E**TGIPIYEAEPLLLFFILFTLLGAIGALGDRGKFVDDPPTGLEKAVIPPGKNVRSALGLKEQGPLFGFTKANELFVGRLAQLGIAFSLIGEIITGKGALAQLNI**E**TGIPIQDIEPLVLLNVAFFFFAAINPGNGKFITDDGEES

Chloroplast transit peptide cleavage site (highlighted with red color):

Cleavage sites residue position: 52-53

The cleavage site is between these two residues.

The cleavage site is the same as reported in Li et al. JBC, 2004 (Regulation of Photosynthetic Light Harvesting Involves Intrathylakoid Lumen pH Sensing by the PsbS Protein).

**E, conserved glutamate residues, to sense thylakoid lumen pH**

AlphaFold2 confidence score (0~100): 76.81662387722845

========

Sevir.5G400800, SvPsbS

Sequence:

MAPSMLMSTSVSGGRALPSLQAARPAAAYPRLALPSVNRHSKSVSVKTLALFGKSKAAKAAPAKKVAAPKPKVEDGIFGTSGGIGFTKENELFVGRVAMLGFAASLLGEAITGKGILAQLNL**E**TGIPIYEAEPLLLFFILFTLLGAIGALGDRGTFVDDVTGLDKAVIQPGKGFRGALGLSEGGPLFGFTKSNELFVGRLAQLGVAFSIIGEIITGKGALAQLNI**E**TGVPINEIEPLVLFNVLFFFIAAINPGTGKFIIGDDEKE

Chloroplast transit peptide cleavage site (highlighted with red color):

Cleavage sites residue position: 57-58

The cleavage site is between these two residues.

**E, conserved glutamate residues, to sense thylakoid lumen pH**

AlphaFold2 confidence score (0~100): 78.60569773719035

========

AT5G67030, AtZEP

Sequence:

MGSTPFCYSINPSPSKLDFTRTHVFSPVSKQFYLDLSSFSGKPGGVSGFRSRRALLGVKAATALVEKEEKREAVTEKKKKSRVLVAGGGIGGLVFALAAKKKGFDVLVFEKDLSAIRGEGKYRGPIQIQSNALAALEAIDIEVAEQVMEAGCITGDRINGLVDGISGTWYVKFDTFTPAASRGLPVTRVISRMTLQQILARAVGEDVIRNESNVVDFEDSGDKVTVVLENGQRYEGDLLVGADGIWSKVRNNLFGRSEATYSGYTCYTGIADFIPADIESVGYRVFLGHKQYFVSSDVGGGKMQWYAFHEEPAGGADAPNGMKKRLFEIFDGWCDNVLDLLHATEEEAILRRDIYDRSPGFTWGKGRVTLLGDSIHAMQPNMGQGGCMAIEDSFQLALELDEAWKQSVETTTPVDVVSSLKRYEESRRLRVAIIHAMARMAAIMASTYKAYLGVGLGPLSFLTKFRVPHPGRVGGRFFVDIAMPSMLDWVLGGNSEKLQGRPPSCRLTDKADDRLREWFEDDDALERTIKGEWYLIPHGDDCCVSETLCLTKDEDQPCIVGSEPDQDFPGMRIVIPSSQVSKMHARVIYKDGAFFLMDLRSEHGTYVTDNEGRRYRATPNFPARFRSSDIIEFGSDKKAAFRVKVIRKTPKSTRKNESNNDKLLQTA

Chloroplast transit peptide cleavage site (highlighted with red color):

Cleavage sites residue position: 60-61

The cleavage site is between these two residues.

AlphaFold2 confidence score (0~100): 85.7710608585738

========

Sevir.7G124500, SvZEP1

Sequence:

MTSTMPSSRTSISLPFASPATRRGSRTLRLLALPTPPAPRRRLGLPPGRARRPVSAVASAAMPAPGPKARVLVAGGGIGGLVFALAAQRKGFEVLVLERDMSAIRGEGRYRGPIQLQSNALAVLEAVDAAAADEVMDAGCVTGDRVNGIVDGISGSWYCKFDTFTPAAERGLPVTRVISRMTLQQILARAVGDDAILNGSHVVDFIDDGSKVTAILEDGRRFEGDLLVGADGIWSKVRKTLFGHSEATYSGYTCYTGIADFVPPDIDTVGYRVFLGHKQYFVSSDVGAGKMQWYAFHKEEAGGTDPENGKKKRLLEIFSGWCDNVIDLINATEEEAILRRDIYDRPPTINWGKGRVTLLGDSVHAMQPNLGQGGCMAIEDGYQLAVELENAWQESVKSGTPMDIVSSLKRYEKERRLRVAIIHGLARMAAIMATTYRPYLGVGLGPLSFLTKLRIPHPGRVGGRFFIMIGMPAMLSWVLGGNSSKLEGRPLSCRLSDKANDQLYRWFEDDDALEQAMGGEWYLFPISEGNSNSLQPVRLIRDEQRAISFGNRSDPSDSASSLALPMPQISERHATITCKNKAFYLTDLGSEHGTWITDNEGRRYRVPPNYPVRFHPSDVIEFGSDQKAMFRVKVLNTLPYESARRGKQQQQQQVLQAA

Chloroplast transit peptide cleavage site (highlighted with red color):

Cleavage sites residue position: 58-59

The cleavage site is between these two residues.

AlphaFold2 confidence score (0~100): 85.15429022255908

========

AT1G08550, AtVDE

Sequence:

MAVATHCFTSPCHDRIRFFSSDDGIGRLGITRKRINGTFLLKILPPIQSADLRTTGGRSSRPLSAFRSGFSKGIFDIVPLPSKNELKELTAPLLLKLVGVLACAFLIVPSADAVDALKTCACLLKGCRIELAKCIANPACAANVACLQTCNNRPDETECQIKCGDLFENSVVDEFNECAVSRKKCVPRKSDLGEFPAPDPSVLVQNFNISDFNGKWYITSGLNPTFDAFDCQLHEFHTEGDNKLVGNISWRIKTLDSGFFTRSAVQKFVQDPNQPGVLYNHDNEYLHYQDDWYILSSKIENKPEDYIFVYYRGRNDAWDGYGGAVVYTRSSVLPNSIIPELEKAAKSIGRDFSTFIRTDNTCGPEPALVERIEKTVEEGERIIVKEVEEIEEEVEKEVEKVGRTEMTLFQRLAEGFNELKQDEENFVRELSKEEMEFLDEIKMEASEVEKLFGKALPIRKVR

Chloroplast transit peptide cleavage site (highlighted with red color):

Cleavage sites residue position: 113-114

The cleavage site is between these two residues.

AlphaFold2 confidence score (0~100): 74.6947939224433

========

Sevir7G073400, SvVDE

Sequence:

MMSRHCANRVFPPGGSNILHGPKSRARAATGRHSAVRFRRCCVRANFWRSDHLPLKVTPSEIIEVLQASDVFGSVKKWSRLQLVTMTGLVACVVLVVPSADAVDALKTCTCLLKECRIELAKCIANPSCAANVACLNTCNNRPDETECQIKCGDLFENSVVDEFNECAVSRKKCVPRKSDVGEFPVPDPSALVKNFNMADFNGKWYISSGLNPTFDTFDCQLHEFHVEGDKLIANITWRIRTPDSGFFTRSTVQRFVQDPSQPGILYNHDNEFLHYQDDWYIISSKVENKDDDYIFVYYRGRNDAWDGYGGSVLYTRSKTVPETIIPELEKAAKSVGRDFSTFIRTDNTCGPEPPLVERIEKTVEEGEKTIIKEVKEIEGEIEEEVEELEKEEVTLFQKLAEGLIEVKQDLMNFLQGLSKEEMELLDQMNMEATEVEKVFSRSLPLRKLR

Chloroplast transit peptide cleavage site (highlighted with red color):

Cleavage sites residue position: 102-103

The cleavage site is between these two residues.

AlphaFold2 confidence score (0~100): 76.81662387722845

========

**Part 2:**

**Predicted interaction sites for Sv/AtPsbS, to support Figure 2**

AT1G44575, AtPsbS

MAQTMLLTSGVTAGHFLRNKSPLAQPKVHHLFLSGNSPVALPSRRQSFVPLALFKPKTKAAPKKVEKPKSKVEDGIFGTSGGIGFTKANELFVGRVAMIGFAASLLGEALTGKGILAQLNL**E**TGIPIYEAEPLLLFFILFTLLGAIGALGDRGKFVDDPPTGLEKAVIPPGKNVRSALGLKEQGPLFGFTKANELFVGRLAQLGIAFSLIGEIITGKGALAQLNI**E**TGIPIQDIEPLVLLNVAFFFFAAINPGNGKFITDDGEES

Sevir.5G400800, SvPsbS

MAPSMLMSTSVSGGRALPSLQAARPAAAYPRLALPSVNRHSKSVSVKTLALFGKSKAAKAAPAKKVAAPKPKVEDGIFGTSGGIGFTKENELFVGRVAMLGFAASLLGEAITGKGILAQLNL**E**TGIPIYEAEPLLLFFILFTLLGAIGALGDRGTFVDDVTGLDKAVIQPGKGFRGALGLSEGGPLFGFTKSNELFVGRLAQLGVAFSIIGEIITGKGALAQLNI**E**TGVPINEIEPLVLFNVLFFFIAAINPGTGKFIIGDDEKE

Interacted residues: 11 sites

| **AtPsbS** | **SvPsbS** |
| --- | --- |
| 71-73 | 156-158 |
| 76-77 | 177 |
| 77-78 | 192-193 |
| 79-81 | 171-173 |
| 126-128 | 226-227 |
| 145-146 | 146-147 |
| 155-157 | 72-74 |
| 171-172 | 80-82 |
| 177 | 78 |
| 193 | 78-79 |
| 226-227 | 127-129 |

AlphaFold-Multimer confidence score (0~1): 0.70027759887

Purple color highlights the sites near or including the conserved pH-sensing glutamate residues.

**Predicted interaction sites for At/ AtPsbS, to support supplemental Figure S3b**

AT1G44575, AtPsbS

MAQTMLLTSGVTAGHFLRNKSPLAQPKVHHLFLSGNSPVALPSRRQSFVPLALFKPKTKAAPKKVEKPKSKVEDGIFGTSGGIGFTKANELFVGRVAMIGFAASLLGEALTGKGILAQLNL**E**TGIPIYEAEPLLLFFILFTLLGAIGALGDRGKFVDDPPTGLEKAVIPPGKNVRSALGLKEQGPLFGFTKANELFVGRLAQLGIAFSLIGEIITGKGALAQLNI**E**TGIPIQDIEPLVLLNVAFFFFAAINPGNGKFITDDGEES

Interacted residues: 9 sites

| **AtPsbS** | **AtPsbS** |
| --- | --- |
| 70-73 | 155-159 |
| 77-78 | 192-193 |
| 78-79 | 179 |
| 226-227 | 126-128 |
| 145-146 | 145-146 |
| 155-159 | 70-73 |
| 179 | 78-79 |
| 192-193 | 77-78 |
| 226-227 | 126-128 |

AlphaFold-Multimer confidence score (0~1): 0.705731046

**Predicted interaction sites for Sv/SvPsbS, to support supplemental Figure S3c**

Sevir.5G400800, SvPsbS

MAPSMLMSTSVSGGRALPSLQAARPAAAYPRLALPSVNRHSKSVSVKTLALFGKSKAAKAAPAKKVAAPKPKVEDGIFGTSGGIGFTKENELFVGRVAMLGFAASLLGEAITGKGILAQLNL**E**TGIPIYEAEPLLLFFILFTLLGAIGALGDRGTFVDDVTGLDKAVIQPGKGFRGALGLSEGGPLFGFTKSNELFVGRLAQLGVAFSIIGEIITGKGALAQLNI**E**TGVPINEIEPLVLFNVLFFFIAAINPGTGKFIIGDDEKE

Interacted residues: 7 sites

| **SvPsbS** | **SvPsbS** |
| --- | --- |
| 71-74 | 156-158 |
| 78-79 | 192-193 |
| 127-129 | 226-227 |
| 146-147 | 146-147 |
| 156-158 | 71-74 |
| 192-193 | 78-79 |
| 226-227 | 127-129 |

AlphaFold-Multimer confidence score (0~1): 0.69361217
